# Phototropin localization and interactions regulates photophysiological processes in *Chlamydomonas reinhardtii*

**DOI:** 10.1101/2024.12.27.630506

**Authors:** Sunita Sharma, Kumari Sushmita, Rajani Singh, Sibaji K. Sanyal, Suneel Kateriya

## Abstract

Phototropin, a blue-light sensing serine/threonine kinase, plays a pivotal role in regulating diverse photophysiological processes in both plants and algae. In *Chlamydomonas reinhardtii*, phototropin (CrPhot) localizes to the eyespot and flagella, coordinating key cellular functions such as phototaxis, photosynthesis, gametogenesis, and chlorophyll biosynthesis. While previous research has identified phototropin interactions with signaling proteins such as channelrhodopsins and light-harvesting complex proteins, many aspects of its interaction network and regulatory mechanisms remain unresolved. In this study, we explored novel interacting protein partners of phototropin and their roles in modulating its regulatory functions in *Chlamydomonas reinhardtii*. Employing a suite of intraflagellar transport (IFT) mutants of *C. reinhardtii* such as IFT172, IFT52, IFT88, IFT139, kinesin/dynein, CEP290 etc., we elucidate that phototropin localization within the flagella and eyespot is IFT-mediated. Our study highlights interaction of phototropin with other photoreceptors-channelrhodopsins (ChR1 and ChR2), chlamyopsin 6, LOV-histidine kinases (LOV-HK1, LOV-HK2) and signaling protein-14-3-3. CRISPR-Cas9 knockouts of phototropin showed reduced ChR1, 14- 3-3 levels and exhibited impaired photomotility. Moreover, two LOV-domain containing histidine kinases, LOV-HK1 and LOV-HK2, were identified in *C. reinhardtii*. Gene expression of LOV-HK1 and LOV-HK2 were found to be elevated in UV-light in *C. reinhardtii* and their genes expression was found to be altered in phototropin CRISPR-Cas9 knockouts. This study provides new insights into phototropin signalosome and highlights molecular mechanisms governing its function. The research outcomes advances our understanding of phototropin trafficking and signal modulation in *Chlamydomonas reinhardtii*, and sets the stage for further exploration into the broader physiological roles of phototropin in cellular responses.

**Graphical abstract:** Phototropin, a blue-light receptor in *Chlamydomonas reinhardtii*, localizes to the flagella and eyespot, mediates phototaxis and photosynthesis. Its trafficking is mediated by intraflagellar transport (IFT) machinery, with mutations in IFT components (kinesin, dynein, IFT172, IFT52, IFT88, IFT139, CEP290) disrupting phototropin localization. Phototropin interacts with other photoreceptors like channelrhodospins (ChR1/2), chlamyopsin 6, LOV-histidine kinases (LOV-HK1, LOV-HK2) and signaling proteins (14-3-3), coordinating light-driven responses. These findings underscore the details of phototropin trafficking and phototropin signaling impacting light-induced physiological processes in *C. reinhardtii*.

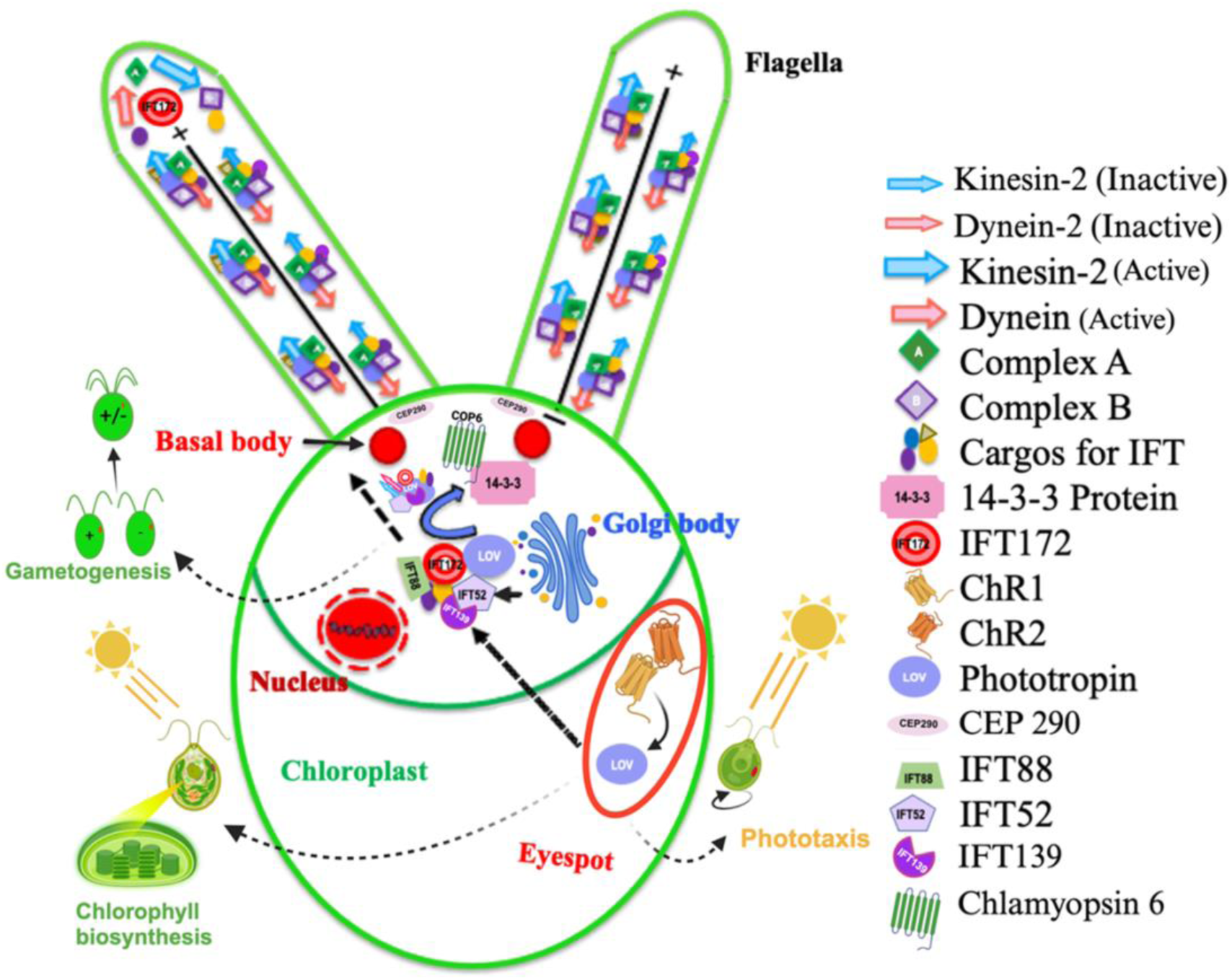

**Highlights:** • Phototropin localizes in eyepot and flagella in *Chlamydomonas reinhardtii*.

• Intraflagellar transport (IFT) mutants of *C. reinhardtii* suggest role of different IFT proteins in phototropin trafficking and localization.

• Phototropin interacts with other photoreceptors (ChR1 & ChR2, COP6, LOV-HK1 & LOV-HK2) and signaling proteins (14-3-3), contributing to various physiological processes.

• CRISPR-Cas9 knockouts of phototropin showed reduced 14-3-3 protein content and photomotility response in *C. reinhardtii*.

## Introduction

Phototropin, a blue-light sensing LOV domain-coupled serine/threonine kinase, orchestrates a range of photophysiological functions in plants and algae. Comprising two LOV domains at the N-terminus and a serine/threonine kinase domain at the C-terminus, phototropin transitions between an inactive and active state. Inactive state occurs in the dark-with a noncovalent linkage between the LOV domain (LOV447) and FMN chromophore, and an active state upon blue light exposure, which establishes a covalent cysteine adduct between the conserved cysteine of the LOV domain (LOV390) and the FMN (C4a) atom [1]. This photoactivation underlies diverse processes such as plant growth promotion, inhibition of hypocotyl elongation, chloroplast rearrangement, leaf movement, and stomatal opening in higher plants [2].

*Chlamydomonas reinhardtii*, with its phototropin (CrPhot), provides a robust model for exploring photoreceptor roles in phototaxis, eyespot development [3], gametogenesis [4], photosynthesis [5], chlorophyll biosynthesis [6], completion of the sexual life cycle [4], chemotaxis towards ammonium ions [7] and photoprotection [2]. Notably, phototropin in *C. reinhardtii* is located both in the eyespot and flagella, suggesting a complex role in cellular signaling and environmental responses. It was found that phototropin trafficking is mediated by intraflagellar transport (IFT) machinery in *C. reinhardtii*. Intraflagellar transport machinery (IFT) is a highly organised motor-driven bidirectional protein transportation machinery in cilia and flagella. Ciliopathies, a group of disorders arising from genetic mutations, result in defective proteins that disrupt cilia formation or function, often due to abnormalities in the IFT process. IFT was first identified in *Chlamydomonas* and serves as an excellent model organism to study IFT. Previous studies have recognized the role of intraflagellar transport (IFT) machinery in the trafficking of phototropin [8]; however, the intricacies of this interaction remain poorly understood.

Our research aims to dissect the molecular mechanisms underlying the role of IFT machinery in phototropin trafficking within *C. reinhardtii*. Using various IFT mutants, we investigated how specific IFT components influence the localization and dynamics of phototropin. Distinct cellular localization of phototropin within eyespot and flagella suggest variations in its specific physiological roles and the involvement of IFT machinery within the cell. It is known that phototropin regulate different physiological processes through interaction with other signaling proteins like channelrhodopsins (ChR1 & ChR2) [3], light harvest complex proteins [5] & other signaling proteins which are not well known. Thus, this study also investigates the novel interacting partners of phototropin and their roles in regulating these diverse functions.

Utilizing CRISPR-Cas9 knockouts of phototropin, we examined the physiological impacts and identified key interactors that influences cellular regulation of phototropin. Our findings are expected to uncover molecular mechanisms governing phototropin function in *Chlamydomonas reinhardtii*, with a focus on its trafficking, interactions with other photoreceptors, and its role in regulating photophysiological processes.

## Material & methods Bioinformatic analysis

Gene sequences of phototropin and other LOV domain-containing proteins in *Chlamydomonas reinhardtii* were sourced from the Phytozome database (version 12; <u>Phytozome</u>) listed in Table S1. Prediction of conserved domains was conducted using the SMART (http://smart.embl-heidelberg.de/) and NCBI CD-search (https://www.ncbi.nlm.nih.gov/Structure/cdd/wrpsb.cgi) tools. The predicted protein-protein interactions were identified using STRING database (https://string-db.org/), based on experimental data and text mining (Fig. S1). The predicted network was further refined, visualized, and analyzed using Cytoscape 3.7.2 employing metrics such as betweenness centrality, clustering coefficient, and edge betweenness. Visualization parameters included node size for clustering coefficient, node color for betweenness centrality (scale: red to blue), and stroke color for edge betweenness centrality. Gene expression data were retrieved from ChlamyNET (http://viridiplantae.ibvf.csic.es/ChlamyNet/scripts/fpkm_data.txt) and analyzed using the Pheatmap and heatmaply packages in R Studio 4.0.

### *Chlamydomonas* strains and culture conditions

*Chlamydomonas* strains were sourced from the Chlamydomonas Resource Center, University of Minnesota (<u>Chlamy Collection</u>). Wild-type strains were cultured in Tris-Acetate-Phosphate (TAP) media, while mutants were grown in TAP media supplemented with 0.1 g/L arginine. Cultures were maintained at 22°C with a 14-hour light/10-hour dark photoperiod with continuous shaking at 110 rpm. Temperature-sensitive IFT mutants were subjected to heat shock at 33°C for 30, 60, or 90 minutes for immunolocalization studies.

For gametogenesis, wild-type strains of both mating types (+ and -) were cultured until reaching mid-exponential phase (10^7 cells/mL). Cells were resuspended in 10 mM HEPES buffer (pH 7.2), and starved overnight in low light (∼25 μmol photons/m^2/sec) with continuous aeration. Mating was induced by mixing equal volumes of each mating type, achieving cell fusion within one hour.

### Cell lysate preparation and immunoblotting

Cultures were grown to the log phase (O.D._700_ around 0.6–0.7), harvested, and subsequently resuspended in PBS (containing 0.1% protease inhibitor cocktail). Cells were lysed using sonication at 30% amplitude with pulses of 8 seconds On/Off for 20 cycles.

Lysates (total cell lysate, TCL) were clarified by centrifugation at 13,000 rpm for 15 minutes, and the supernatant (soluble fraction, SF) was treated with 2X Laemmli buffer and heated at 65°C for 30 minutes. Proteins were resolved by SDS-PAGE (12%) and immunoblotting was performed using specific antibody against *Chlamydomonas* Phot, 14-3-3 or ChR1, according to the protocol established earlier [23].

### Immunolocalization analysis

For immunolocalization, coverslips were coated with 0.01 mg/mL Poly-L-Lysine (Sigma, USA) and air-dried. *Chlamydomonas* cells were seeded on these coverslips at room temperature for 15 minutes. Fixation was achieved using 3.7% paraformaldehyde in 1X PBS for 4 minutes, followed by permeabilization with ethanol. Cells then underwent for further incubation in 1X PBS containing 250 mM NaCl, and were washed twice with 1X PBS supplemented with 0.5% Triton X-100 (PBST). Overnight incubation at 4°C was performed with primary antibodies against CrPhot (1:250) and acetylated tubulin (1:1000). After four washes with 1X PBST, cells were incubated with fluorophore-conjugated secondary antibodies: anti-goat Alexa-488 and anti-mouse Alexa-647 (1:1000, Invitrogen, USA) for 2 hour at room temperature. Following two final washes with 1X PBST, the specimens were mounted using SlowFade® Gold antifade reagent (Molecular Probes, Invitrogen, USA) and examined with an Olympus Fluoview 1000 confocal microscope [23]. Strains used for the study are mentioned in Table S2.

### RNA isolation, quantitative RT-PCR and gene expression analysis

For gene expression analysis of LOV domain-associated signaling proteins, *Chlamydomonas reinhardtii* strains, including wild-type (CC125) and CRISPR-Cas mutants, were cultured under a 10 hours dark/14hourslight cycle at 22°C until reaching mid-exponential phase. Gene expression was compared between dark-adapted (14 hours) and light-adapted (4 hours) samples, followed by blue light exposure (∼400 lux) at different time intervals (30, 60, and 120 minutes) prior to RNA extraction. Samples not exposed to blue light were used as controls.

To investigate the effects of different light intensities on the expression of photoreceptor genes interacting with phototropin, cells were subjected to various light conditions (red, green, blue, UV, and white). RNA was extracted using the TRIzol reagent, followed by phase separation with chloroform and RNA precipitation with isopropanol followed by washing with 70% ethanol. The RNA pellet was air-dried and re-suspended in DEPC-treated water. cDNA synthesis was performed using the AffinityScript cDNA Synthesis Kit (Agilent, USA) according to manufacturer’s protocol and quantitative PCR (qPCR) was conducted using KAPA SYBR Green Master Mix on the Agilent MX3005P qPCR system. Primers used for amplification are listed in Table S3, with the CBLP (Beta subunit-like polypeptide) gene serving as the internal control for normalization [24]. Data were collected from three independent experiments, each performed in triplicate, with error bars indicating the standard deviation (SD). All the experiments have been performed in triplicates.

## Results

### Bioinformatic characterization of LOV domain-containing proteins in *C. reinhardtii*

Using the LOV domain from *Chlamydomonas* phototropin (CrPhot) as a template, we identified three LOV domain-containing proteins in *C. reinhardtii*: LOV serine/threonine kinase (CrPhot), and two LOV histidine kinases (CrLOV-HK1 and CrLOV-HK2). Domain analysis revealed that CrPhot contains two N-terminal LOV domains (LOV1 and LOV2) and a C-terminal serine/threonine kinase domain. The identified LOV histidine kinases, CrLOV- HK1 and CrLOV-HK2, vary in domain architecture; CrLOV-HK1 features a single N-terminal LOV domain followed by a histidine kinase (HisK) domain and a response regulator (REC) domain at the C-terminus, whereas CrLOV-HK2 comprises an N-terminal LOV domain, a HisK domain, and two REC domains (Fig. 1a). We further identified several ciliary targeting signals in phototropin (Table S4), which are likely critical for its intracellular trafficking and functional roles. Additionally, we conducted a comprehensive analysis of post-translational modifications (PTMs) in *C. reinhardtii* for LOV domain proteins like SUMOylation, phosphorylation, acetylation, ubiquitination, and lipid modifications. These modifications are likely to influence the proteins’ subcellular localization, stability, and interaction capabilities, thereby modulating light-mediated signaling pathways in response to environmental cues. Detailed PTM data, including specific positions and peptide sequences, are summarized in Table S5, with sequences of the respective LOV domain proteins (Phot, LOV-HK1, LOV- HK2) provided in the Supplementary file.

**Fig. 1.**
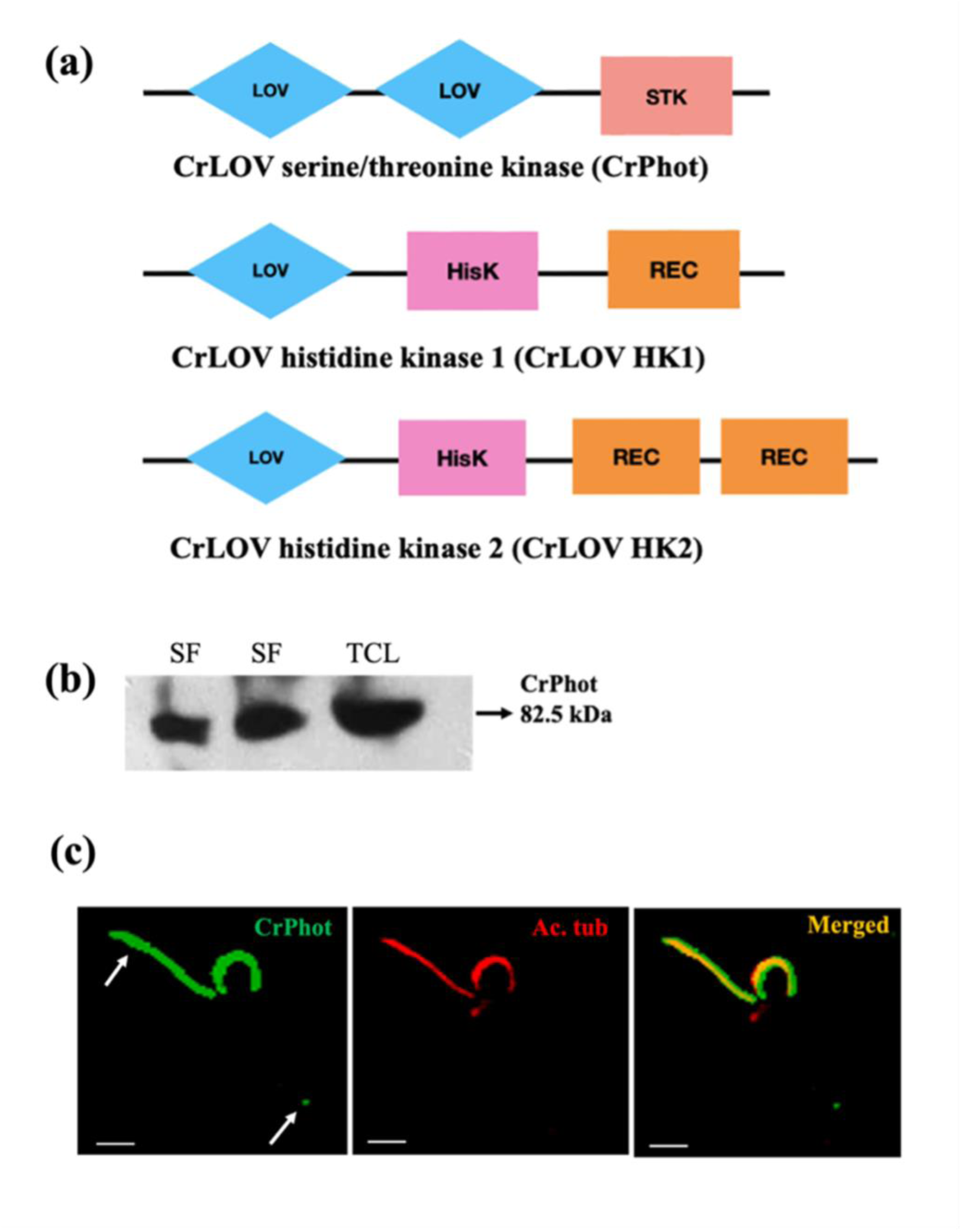
Identification and localization of LOV domain proteins in *Chlamydomonas reinhardtii*. (a) Identification of different type of LOV domain containing proteins in *Chlamydomonas reinhardtii.* Three LOV domain containing proteins LOV serine/threonine kinase (CrPhot), LOV histidine kinases (CrLOV-HK1 and CrLOV-HK2) were identified from *C. reinhardtii* database. (b) Phototropin (CrPhot) immunoblot from TCL (*Chlamydomonas reinhardtii* total cell lysate) with anti-cLOV (with dilution 1:1000) antibody; (c) Immunolocalization of CrPhot in eyespot and flagella with anti-cLOV(green) and anti-acetylated tubulin (red) antibody. Arrow indicates the localization of CrPhot in eyespot and flagella in cell. Scale bar = 2 μm; act. tubulin: acetylated tubulin; LOV: Light-Oxygen-Voltage; NSP: Nitrile specifier protein; STK: Serine/threonine kinase.

### Cellular detection of phototropin in *Chlamydomonas reinhardtii* (CrPhot)

Immunodetection of phototropin with antibody against its LOV domain (anti-cLOV) resulted in full-length phototropin in the cellular fraction at an apparent molecular weight of 82.5 kDa (Fig. 1b). Immunolocalization in the wild type strain (CC-125) demonstrated that at a given condition, CrPhot primarily localizes to the eyespot and flagella of *C. reinhardtii* (Fig. 1c).

### Role of IFT172 in phototropin trafficking at the flagellar tip

IFT172, a component of the intraflagellar transport complex-B, is crucial for the anterograde transport of proteins from the base to the tip of flagella. Precisely, at the flagellar tip, it regulates the cargo transition from anterograde to retrograde transport. The temperature-sensitive flagellar assembly mutant CC-1920 harbors a mutation (L1615P) in the IFT172 gene (*fla11*) [25]. In the present study, at a restrictive temperature of 33°C, CrPhot accumulation was observed at the flagellar tip and near basal body in the IFT172 mutant, suggesting role of IFT172 in phototropin switching at the flagellar tip from anterograde to retrograde machinery [26]. At permissive temperatures, CrPhot was evenly distributed along the flagella, indicating normal IFT function (Fig. 2). These results suggest a significant role of the IFT172 in the cellular trafficking of phototropin within *C. reinhardtii* flagella.

**Fig. 2.**
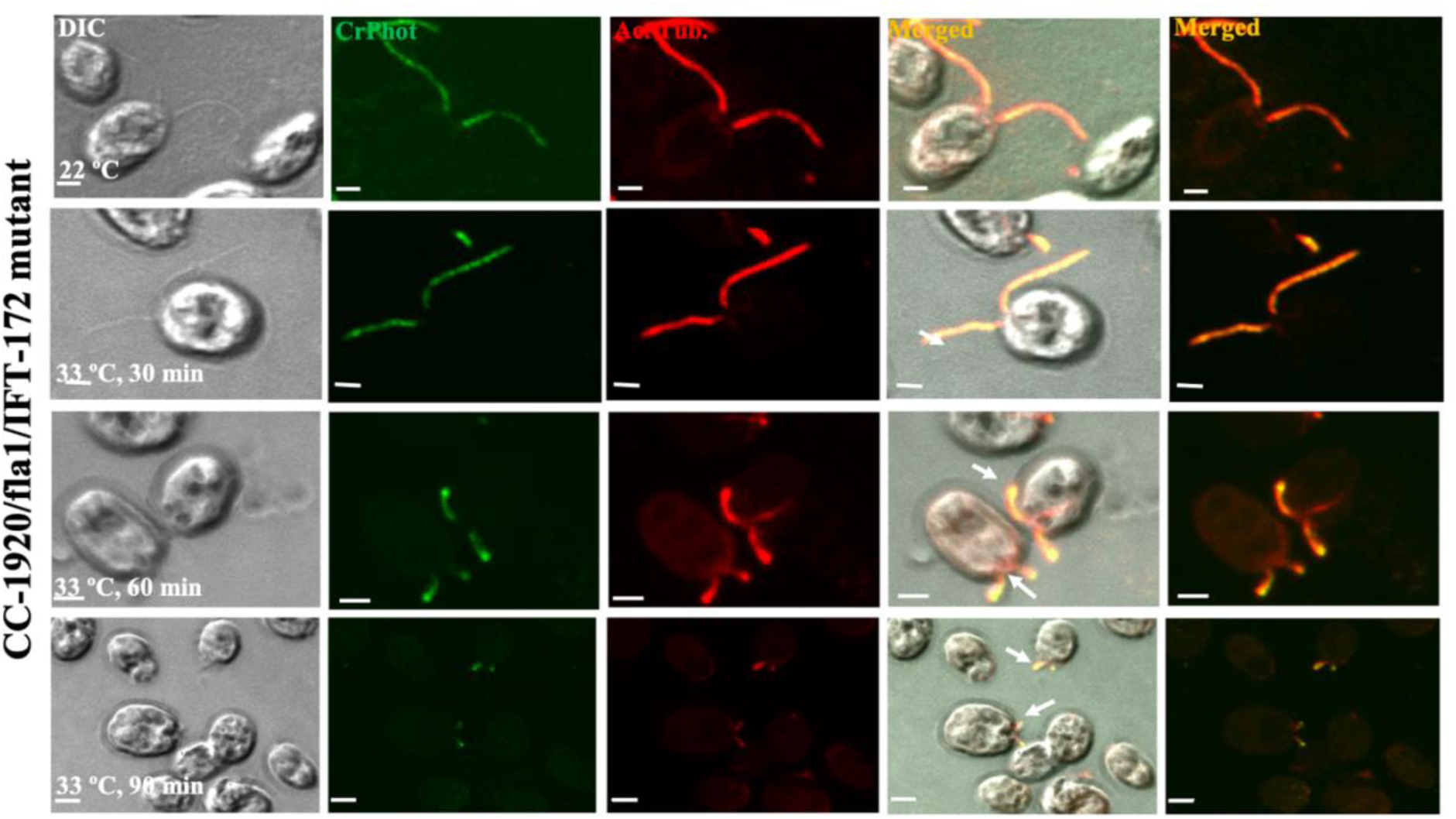
Immunolocalization of CrPhot in IFT172 mutant (CC-1920) strain of *C. reinhardtii*. The accumulation of phototropin was observed at the flagellar tip and near basal body in the mutant strain. The first panel represents DIC image of algal cells. The second panel represents the CrPhot signal (green) with primary antibody anti-cLOV (dilution-1:250) and secondary antibody anti-goat IgG Alexa-488 (dilution-1:1000). The third panel represents the signal for acetylated tubulin (red) with primary antibody anti-act. tubulin (dilution-1:1000) and secondary antibody anti-mouse IgG Alexa-647 conjugated (dilution-1:1000). The fourth and fifth panel represent the overlay (merged) of the first three panels. Scale bar = 2 μm; min: minutes; CrPhot: *Chlamydomonas* phototropin; act. tubulin: acetylated tubulin. The arrow indicates the accumulation of phototropin.

### Impact of IFT52 and IFT88 mutations on anterograde IFT and phototropin stability

Mutations in IFT88 and IFT52, core components of IFT complex-B, critically undermine the stability of the anterograde transport system which is essential for protein trafficking within flagella [27]. IFT52 is essential for assembling the IFT-B core complex and IFT88 stabilizes the IFT-B core complex by directly interacting with IFT52 [27,28]. The IFT52 (CC-477) and IFT88 (CC-3943) mutant strains cultivated at 22°C, revealed significant flagellar abnormalities. These strains displayed either completely absent or markedly shortened flagella as a direct result of defects in IFT52 and IFT88. Immunofluorescence study in CC-477 and CC-3943 strains showed strikingly different result as compared to wild type. In contrast to the wild-type strain CC-124, no detectable CrPhot signal was observed in these mutants (Fig. 3a), despite the presence of consistent protein loading (CrPhot protein) in the cellular lysates as shown by immunoblot analysis (Fig. 3b). The observed degradation of CrPhot suggests that mutations in IFT52 and IFT88 disrupt the stability of the IFT complex, impeding normal phototropin transport consequently, disrupting phototropin stability within cells. These findings emphasize the critical role of IFT52 and IFT88 in maintaining functional phototropin trafficking and its stability in *Chlamydomonas reinhardtii*.

**Fig. 3.**
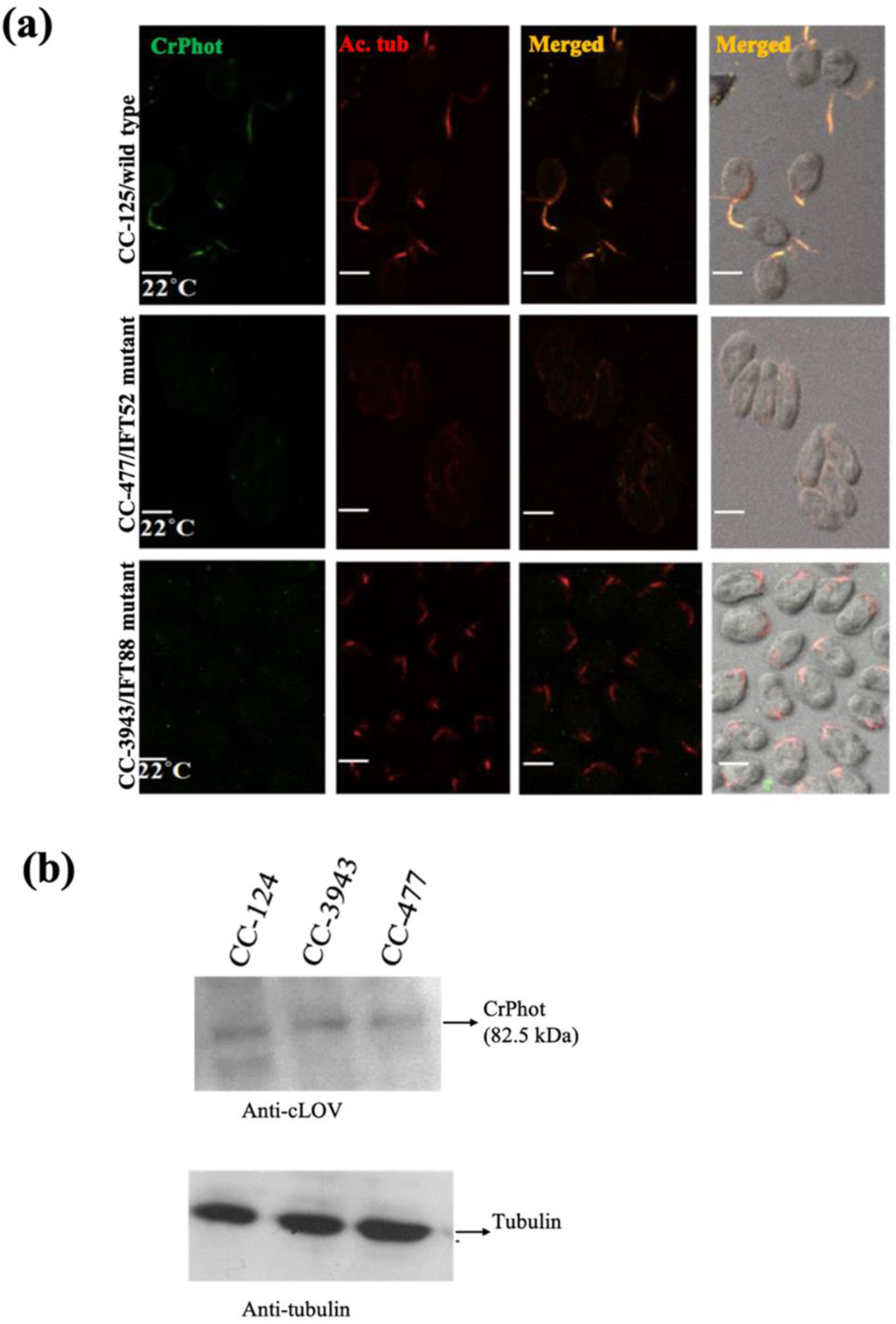
Immunolocalization and expression profile of CrPhot in IFT52 (CC-477) and IFT88 (CC-3943) mutant strains of *C. reinhardtii* cells. Despite normal phototropin synthesis no phototropin signal detected in the mutant strain as compared to wild type. (a) Immunolocalization of CrPhot in IFT52 (CC-477) and IFT88 (CC- 3943) mutant strains with the first panel represents the CrPhot signal (green) with primary antibody anti-cLOV (dilution-1:250) and secondary antibody anti-goat IgG Alexa-488 (dilution-1:1000). The second panel represents the signal for acetylated tubulin (red) with primary antibody anti-act. tubulin (dilution-1:1000) and secondary antibody anti-mouse IgG Alexa-647 conjugated (dilution-1:1000). The third and fourth panel represent the overlay (merged) of the first two panels. Scale bar = 2 μm; min: minutes; CrPhot: *Chlamydomonas* phototropin; act. tubulin: acetylated tubulin. (b) Expression profile of CrPhot in wild type, IFT52, and IFT 88 mutants using immunoblot analyses with the *Chlamydomonas* total cell lysate (CrTCL) of the wild-type strain, IFT52, and IFT88 mutant strain with anti-cLOV antibody (1:1000). α-tubulin was used as loading control.

### Role of IFT139 in retrograde trafficking of phototropin in *Chlamydomonas reinhardtii*

In *Chlamydomonas reinhardtii*, the protein IFT139, a peripheral component of the retrograde intraflagellar transport (IFT) complex, plays a critical role in the localization and trafficking of phototropin. Analysis of the temperature-sensitive mutant strain CC-3863, which harbors a mutation in IFT139, revealed that IFT139 is essential for maintaining proper phototropin distribution within the flagella. At a permissive temperature of 22°C, phototropin is well localized within the flagella of wild-type cells. However, at the restrictive temperature of 33°C, the IFT139 mutation induces aberrant accumulation of phototropin at the flagellar tip and near basal body regions (Fig. 4). This suggests the significant function of the retrograde IFT machinery in modulating phototropin trafficking, thereby influencing photoreceptive signaling pathways in *C. reinhardtii* [29].

**Fig. 4.**
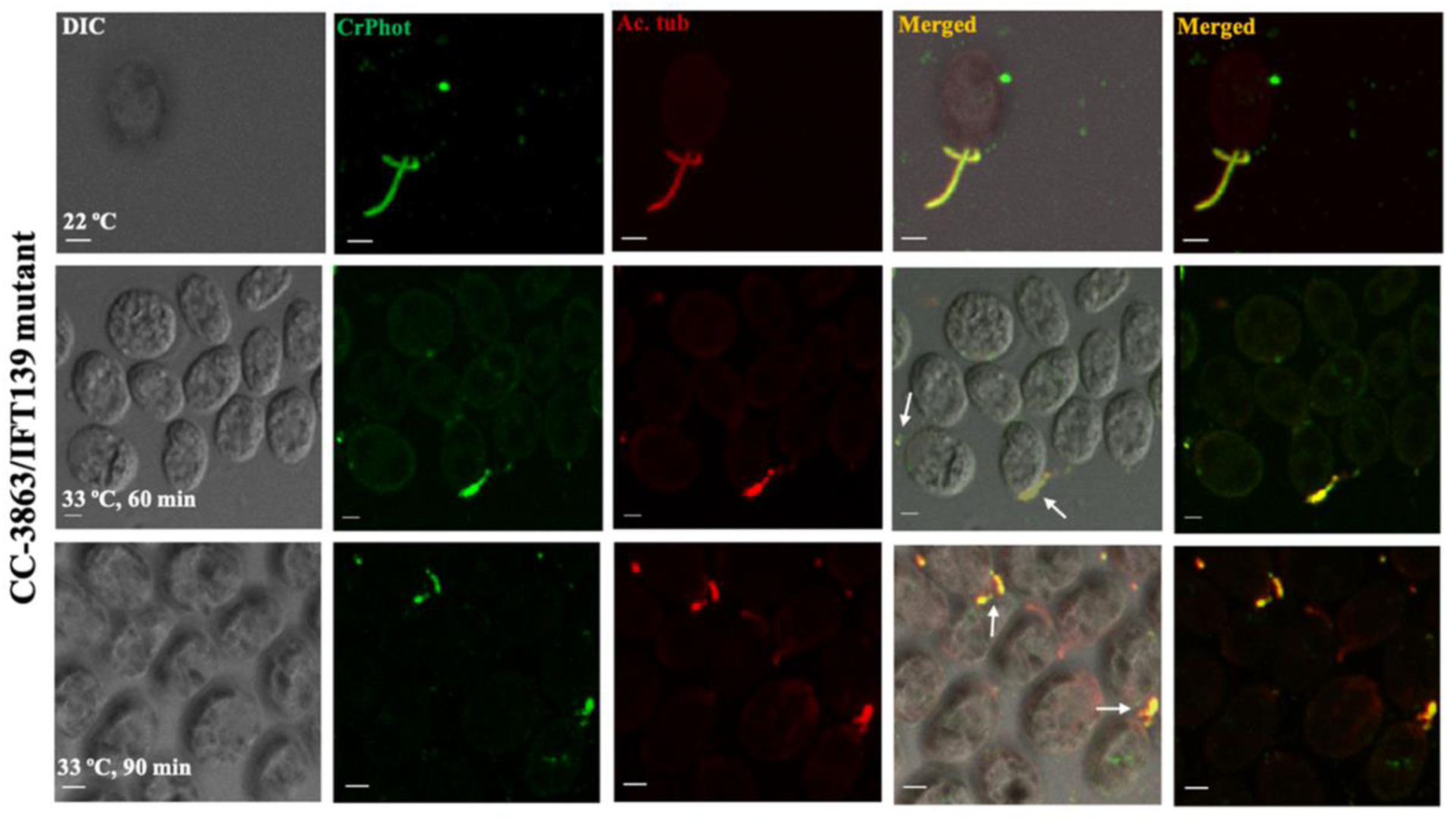
Immunolocalization of CrPhot in IFT139 mutant (CC-3863) strain of *C. reinhardtii* cells. The aberrant accumulation of phototropin was observed at the flagellar tip and near basal body in IFT139 mutant strain. The first panel represents the DIC image of algal cells. The second panel represents the CrPhot signal (green) with primary antibody anti-cLOV (dilution-1:250) and secondary antibody anti-goat IgG Alexa-488 (dilution-1:1000). The third panel represents the signal for acetylated tubulin (red) with primary antibody anti-act. tubulin (dilution-1:1000) and secondary antibody anti-mouse IgG Alexa-647 conjugated (dilution-1:1000). The fourth and fifth panel represent the overlay (merged) of the first three panels. Scale bar = 2 μm; min: minutes; CrPhot: *Chlamydomonas* phototropin; act. tubulin: acetylated tubulin. The arrow indicates the accumulation of phototropin.

### Role of kinesin-2 motor protein (Fla8/Fla10) in phototropin trafficking in *Chlamydomonas reinhardtii*

The anterograde motor protein kinesin plays a critical role in phototropin trafficking within *C. reinhardtii* [30]. We investigated this mechanism using the kinesin-2 motor subunit mutant strain, CC1396 (*fla8*), which harbors a glutamic acid to lysine substitution (E21K) leading to defective assembly and transport of anterograde intraflagellar transport (IFT) particles and cargoes, while retrograde movement remains comparable to the wild type [31,32]. This temperature-sensitive mutant functions like the wild type at permissive conditions (22°C) but exhibits flagellar retraction at non-permissive temperatures (33°C). After 120 minutes of incubation at a non-permissive temperature, CrPhot accumulates at the flagellar tip, contrasting with its localization along the flagella in the wild type strain CC-124 at permissive temperatures, underscoring the kinesin motor’s involvement in phototropin trafficking (Fig. 5a).

**Fig. 5.**
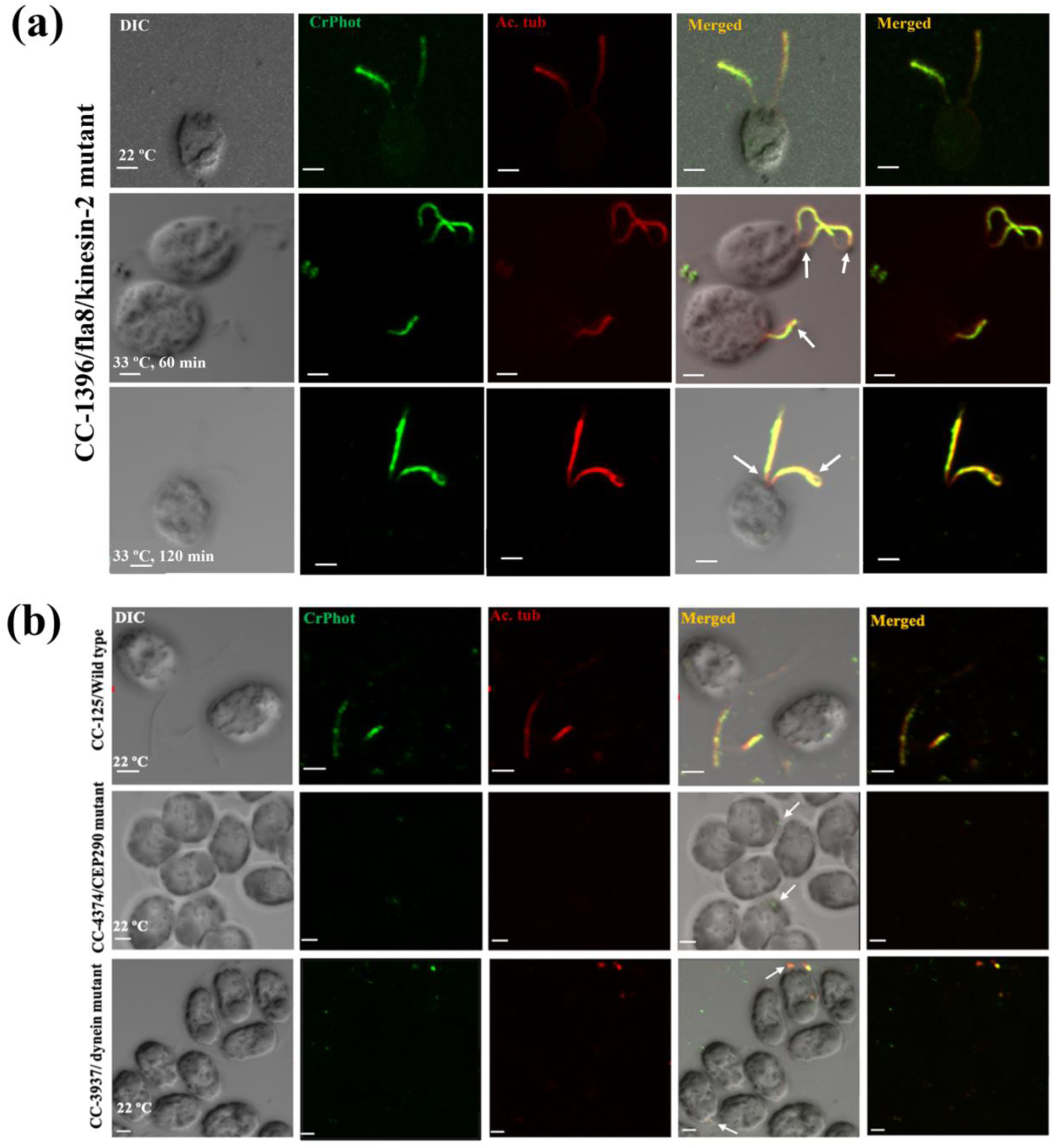
Immunolocalization of CrPhot in kinesin-2 (Fla8) mutant (CC-1396), dynein light chain (LC8) mutant (CC-3937) and centrosome associated protein 290 (CEP290) mutant (CC-4374) strain of *C. reinhardtii*. The accumulation of phototropin at flagellar tip was observed in kinesin and dynein mutant strains, whereas the phototropin signal was absent in CEP290 mutant. The first panel represents the DIC image of algal cells. The second panel represents the CrPhot signal (green) with primary antibody anti-cLOV (dilution-1:250) and secondary antibody anti-goat IgG Alexa-488 (dilution-1:1000). The third panel represents the signal for acetylated tubulin (red) with primary antibody anti-act. tubulin (dilution-1:1000) and secondary antibody anti-mouse IgG Alexa-647 conjugated (dilution-1:1000). The fourth and fifth panel represent the overlay (merged) panels of first three panels. The arrow indicates the accumulation of phototropin. Scale bar = 2 μm; min: minutes; CrPhot: *Chlamydomonas* phototropin; act. tubulin: acetylated tubulin.

Additionally, we examined another kinesin-2 motor subunit mutant, temperature-sensitive strain, CC1919 (*fla10*) [33]. Despite prolonged exposure to non-permissive temperatures (60 and 90 minutes), there was no significant impact on flagellar assembly (Fig. S2); however, we observed protein accumulation at the flagellar tip. These findings provide insight into the mechanistic role of kinesin motors in phototropin trafficking within flagellar assembly pathways of *Chlamydomonas reinhardtii*.

### Role of dynein motor protein (LC8) and centrosome-associated protein (CEP290) in phototropin trafficking in *Chlamydomonas reinhardtii*

In *Chlamydomonas reinhardtii*, the role of IFT machinery in trafficking of phototropin was investigated in a dynein motor protein mutant strain (dynein light chain 8; DLC8), CC-3937. This mutant strain exhibits either absent or significantly truncated flagella even at a permissive temperature of 22°C, attributed to mutations in DLC8. In immunofluorescence study, a marked accumulation of CrPhot at the flagellar tips was observed in the DLC8 mutant (Fig. 5b), highlighting the critical role of the dynein motor protein in phototropin trafficking within the organism.

Further, we studied CrPhot localization in another motor protein mutant strain, CC4424. CC4424 is a temperature-sensitive double mutant defective in the kinesin-2 motor subunit encoded by *fla10* and dynein heavy chain *dhc-1b-3* [34]. Cells in late light phase were used for the experiment and were incubated at non-permissive temperature for different time points (30, 60 and 90 minutes). We found the protein is well localized in flagella at permissive temperature, however, incubation for different time points at non-permissive temperatures (33°C) showed phototropin accumulation at flagellar tip in mutant. It suggests that kinesin or dynein involved in phototropin trafficking in *C. reinhardtii* (Fig. 6).

**Fig. 6.**
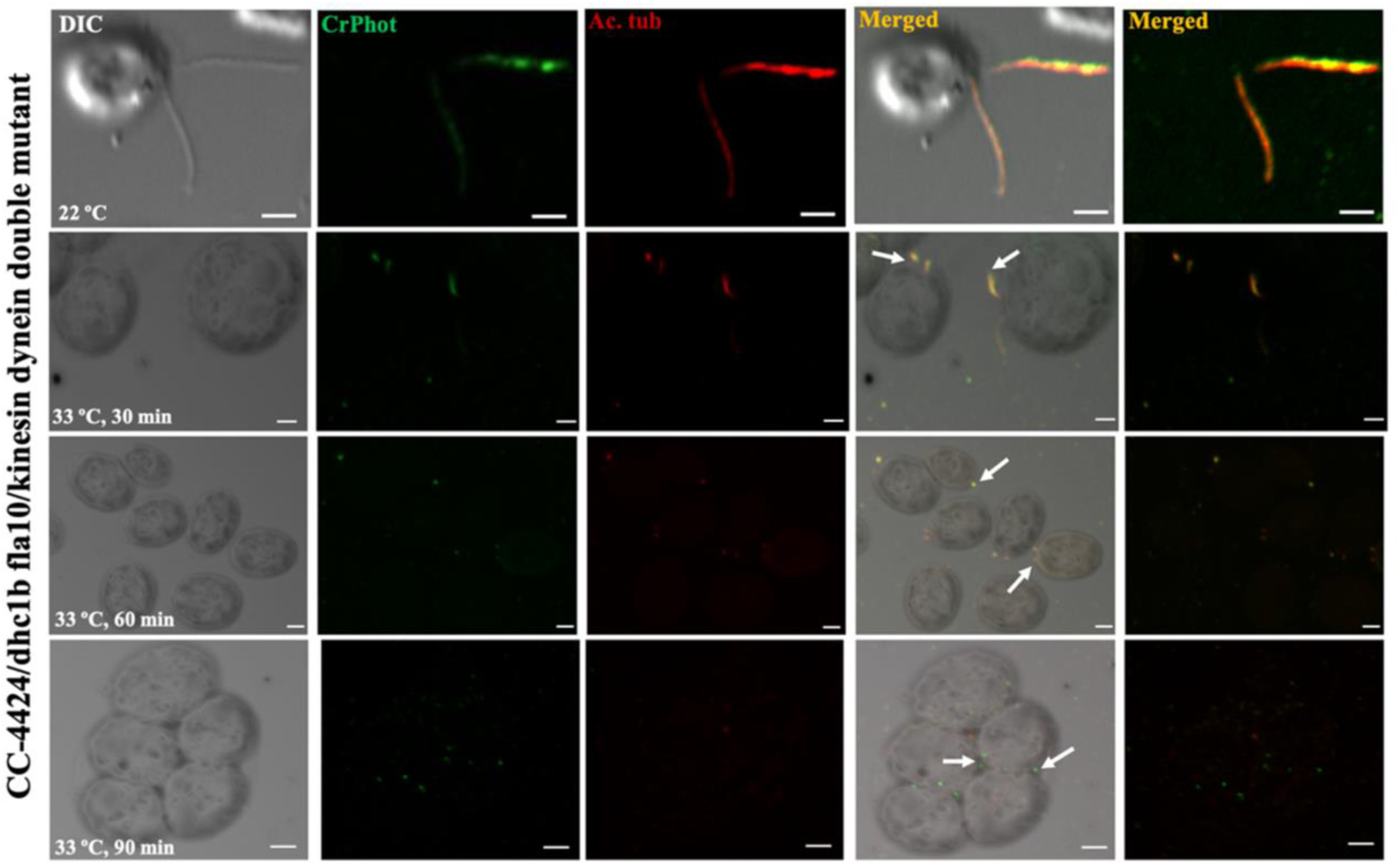
Immunolocalization of CrPhot in dynein heavy chain (Dhc-1b-3) and kinesin-2 (Fla10) double mutant strain (CC-4424) of *C. reinhardtii*. Mutation in both dynein and kinesin results in accumulation of phototropin in double mutant strain of *C. reinhardtii*. The first panel represents the DIC image of algal cells. The second panel represents the CrPhot signal (green) with primary antibody anti-cLOV (dilution-1:250) and secondary antibody anti-goat IgG Alexa-488 (dilution-1:1000). The third panel represents the signal for acetylated tubulin (red) with primary antibody anti-act. tubulin (dilution-1:1000) and secondary antibody anti-mouse IgG Alexa-647 conjugated (dilution-1:1000). The fourth and fifth panel represent the overlay (merged) of the first three panels. The arrow indicates the accumulation of phototropin in kinesin and dynein mutants, while in CEP290 it indicates absence of phototropin signal. Scale bar = 2 μm; min: minutes; CrPhot: *Chlamydomonas* phototropin; act. tubulin: acetylated tubulin.

Another important protein CEP290, a centrosome-associated protein with a molecular mass of 290 kDa, primarily localizes to the centrosomes in actively dividing cells. In *Chlamydomonas reinhardtii*, the absence of the CEP290-containing transition zone results in the accumulation of intraflagellar transport B (IFT-B) subcomplex components within the algal flagella and a reduction in IFT-A subcomplex components [35–37]. Moreover, CEP290 plays a critical role in recruiting BBSome proteins to the flagella [38]. The CC-4374 strain, which harbors mutations in CEP290, exhibits phenotypic defects characterized by extremely short or absent flagella even at a permissive temperature of 22°C. In the CEP290 mutant, the CrPhot signal is notably absent near the basal body (Fig. 5b), a phenomenon likely attributed to the altered dynamics of IFT-B and the diminished presence of IFT-A subcomplex components crucial for protein trafficking. This suggests the critical role of CEP290 in regulating phototropin trafficking and stability, highlighting its essential function in cellular trafficking mechanisms within *C. reinhardtii*.

### Phototropin-mediated crosstalk among different photoreceptors and signaling proteins in *Chlamydomonas reinhardtii*

Phototropin plays a pivotal role in various photophysiological processes, including gametogenesis, photoprotection, photomotility, eyespot development, and completion of the sexual life cycle in *Chlamydomonas reinhardtii*. To identify the interacting protein partners of CrPhot and crosstalk with different photoreceptors curated protein-protein networking was predicted. Interactome depicts primary interaction of phototropin with FTT1 (14-3-3), Cop3 (ChR1), Cop4 (ChR2), LOV-HK1 & LOV-HK2 (LOV-histidine kinases), ADP ribosylation factor-like 3 (ARF), CGL 49, other interacting protein include PINs, calcineurin B-like proteins (CBL), calmodulin (CAM), chlamyopsin 5 (COP5), chlamyopsin 6 (COP6). Additionally, interactome analysis also revealed phototropin association with intraflagellar transport (IFT) components for protein trafficking in the eyespot and flagella (Fig. 7a, Table S6).

**Fig. 7.**
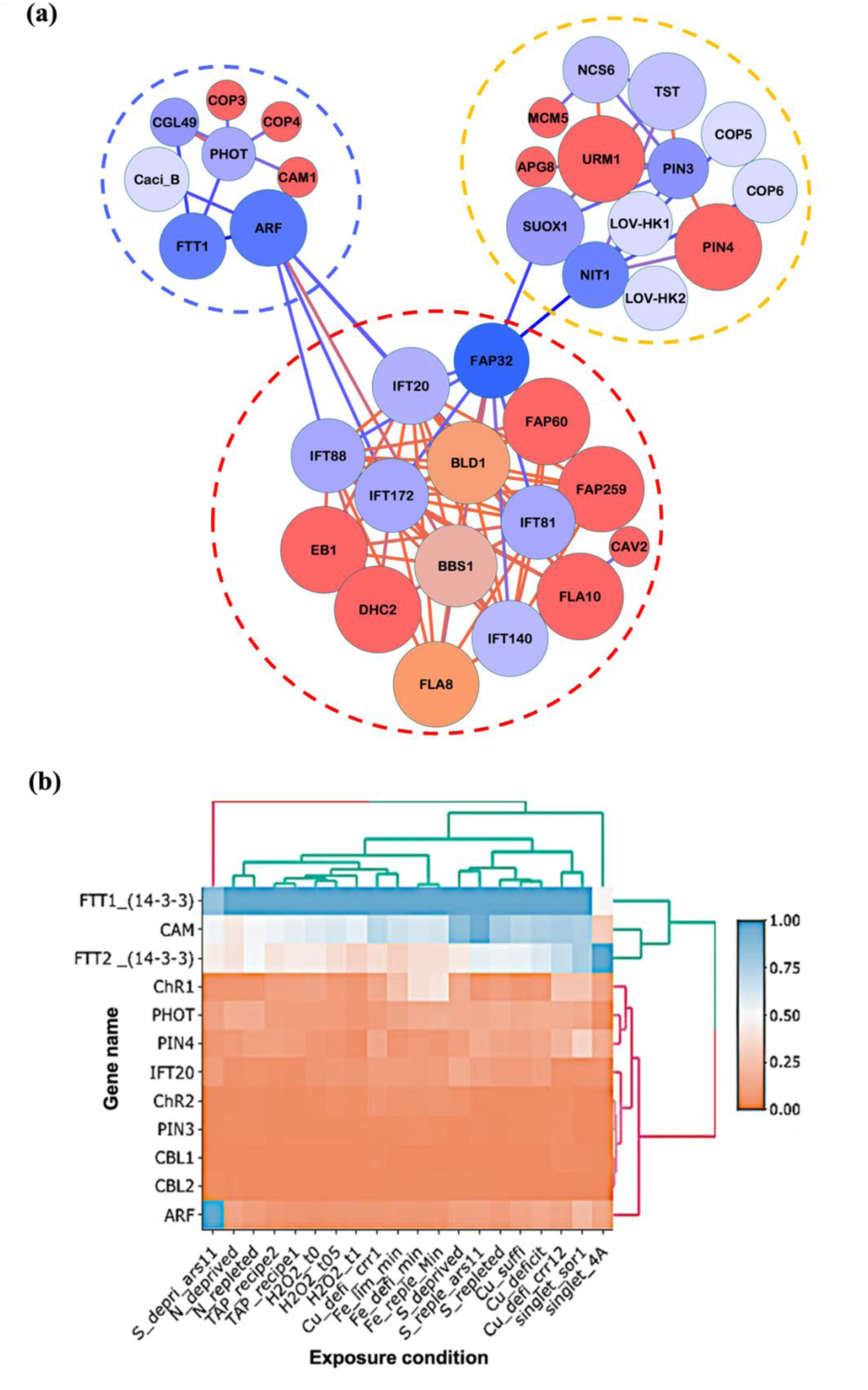
Phototropin-mediated crosstalk among different photoreceptors and signalling proteins in *C. reinhardtii*. Phototropin shows primary interaction with different photoreceptor(s) and 14-3-3. The expression of these genes is regulated by phototropin via specific signalling pathways. (a) Interactome prediction of phototropin (Phot), LOV-histidine kinases (LOV-HK), ADP-ribosylation factor (ARF), rhodopsin (Rh), 14-3-3, calcineurin B-like protein (CBL), intraflagellar transport proteins (IFT), and PIN-FORMED proteins (PIN) in *Chlamydomonas reinhardtii*, generated using STRING software. Key interactions involving PIN, phototropin, and IFT proteins are highlighted by yellow, blue, and red circles, respectively. (b) Expression profile analysis of genes interacting with phototropin in *Chlamydomonas reinhardtii*, retrieved from ChlamyNET and analyzed using R Studio.

To investigate phototropin-mediated signaling in photosynthesis, we analyzed gene expression of predicted phototropin-interacting genes under nutrient depletion and varying light conditions, using ChlamyNET data. Genes in the phototropin interactome, such as IFT20, 14- 3-3, ARF, and calmodulin, showed upregulation, while ChR1, ChR2, PIN3, PIN4, and CBL1/CBL2 were downregulated. This suggests phototropin might regulate these genes through specific signaling pathways (Fig. 7b).

### Interaction of phototropin with 14-3-3 protein and predicted functional roles

The regulatory protein 14-3-3, which is conserved across eukaryotes, is known to bind phosphorylated and non-phosphorylated proteins via specific motif in protein, influencing numerous signaling pathways (Fig. S3) [39]. Based on literature survey, we analyzed photoreceptor and IFT proteins of *C. reinhardtii* for 14-3-3 binding motif. Sequence analysis identified conserved 14-3-3 binding motifs in ChR1, phototropin, COP5, COP6, and several IFT components, including kinesin-2 and IFT172 (Table 1). Sequence alignment of 14-3-3 revealed conserved phosphorylation, dimerization, and calcium-binding motifs underlining its functional conservation in *C. reinhardtii* (Fig. 8d). In *C. reinhardtii*, 14-3-3 has been localized to the eyespot and flagella (Fig. 8a) and detected in cellular fractions (Fig. 8b). Given the established role of phototropin in modulating stomatal opening via 14-3-3 interactions in *Arabidopsis* [40] and its involvement in controlling ChR1 levels in mediating phototaxis [3], immunoblotting was performed on total cell lysates from wild-type and phototropin knockout strains. The absence of phototropin resulted in altered expression of both ChR1 and 14-3-3, with the ΔPhot-B5 and ΔPhot-C4-1 mutants showing reduced levels of these proteins compared to the wild type (Fig. 8c). These findings suggest that phototropin may regulate photomotility and protein expression via a signaling pathway involving 14-3-3 and ChR1, providing further insight into the molecular mechanisms underpinning phototropin-mediated responses in *Chlamydomonas*.

**Fig. 8.**
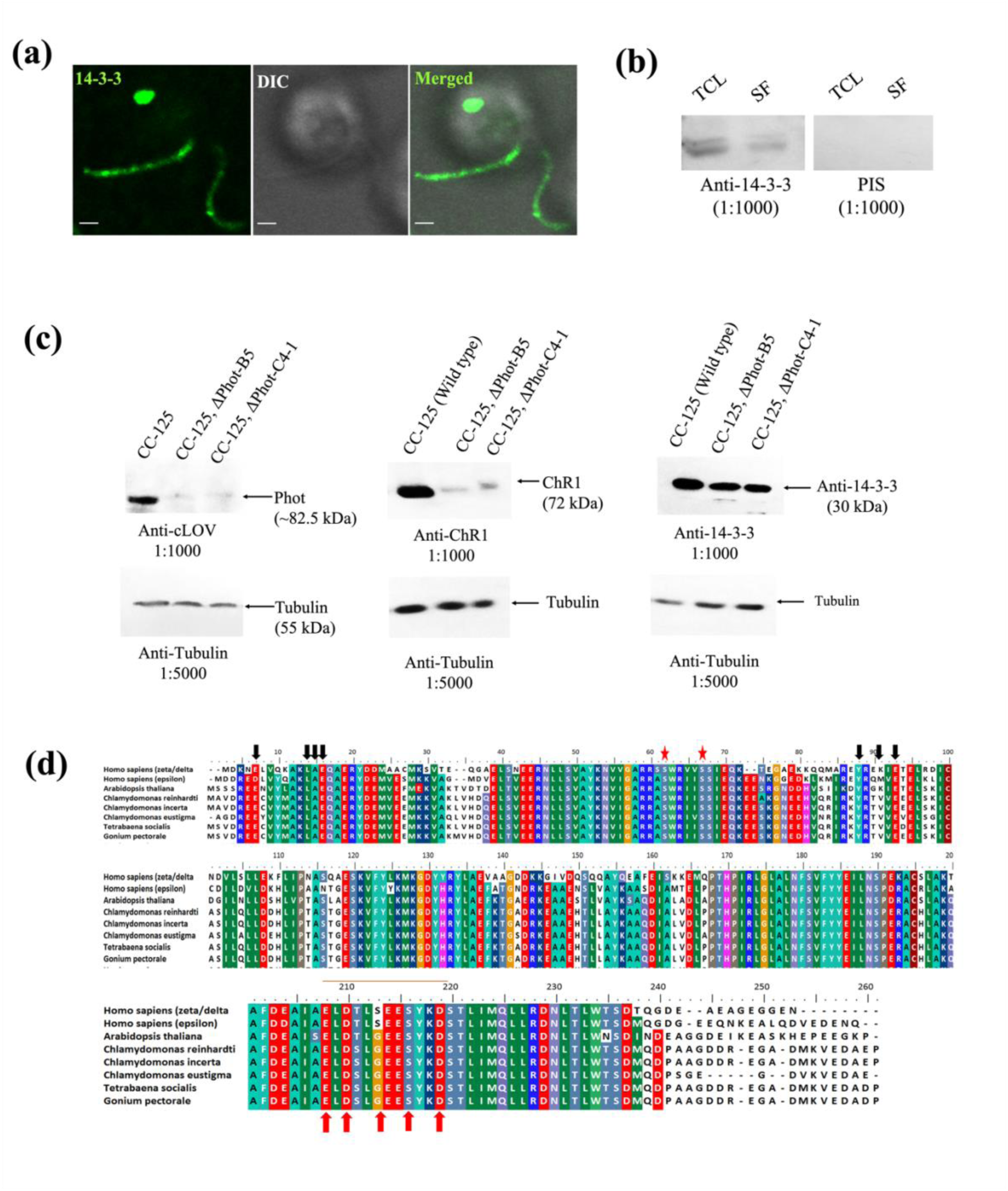
Immunoblotting, immunolocalization, and sequence conservation of 14-3-3 protein in *Chlamydomonas reinhardtii*. Phototropin mutation effect the protein levels of 14-3-3 and ChR1 in phototropin knockouts of *C. reinhardtii*. (a) Immunolocalization of 14-3-3 protein in *Chlamydomonas* algal cells; (b) Immunoblotting of *Chlamydomonas* cell lysate (TCL: total cell lysate, SF: soluble fraction) with anti-14-3-3 and pre- immune serum (PIS) at dilution 1:1000; (c) Expression of ChR1 and 14-3-3 in wild type and phototropin mutant strains of *Chlamydomonas reinhardtii*, as shown by immunoblotting of cLOV1, ChR1, and 14-3-3 in wild type (CC-125) and phototropin knockouts (CC-125, ΔPhot-B5 & CC-125, ΔPhot-C4-1); (d) Sequence alignment of 14-3-3 sequence of *Chlamydomonas reinhardtii* with the sequences from different organisms. Residues shown by black and red arrows represent the conserved amino acids responsible for dimerization in 14-3-3 sequence and EF hand motif characteristic for Ca2+-binding sites, respectively. Residues marked with red stars represent putative phosphorylation sites in the 14- 3-3 sequence. Scale bar = 2 μm.

**Table 1.** Interacting partner of 14-3-3 in *Chlamydomonas*

| **Proteins** | **Form of the binding motif** | **Motif in sequence** |
| --- | --- | --- |
| Phototropin | RX<sub>1-2</sub>pSX<sub>2-3</sub>S and CRD | RANpSFVGT and CRD |
| ChR1 | CRD | CRD |
| COP5 | RX<sub>1-2</sub>pSX<sub>2-3</sub>S | RFMpSSMS |
| COP6 | RX<sub>1-2</sub>pSX<sub>2-3</sub>S | RSLpSGLS |
| Kinesin-2 motor | RX<sub>1-2</sub>pSX<sub>2-3</sub>S | RPVpSAS |
| IFT172 | RX<sub>1-2</sub>pSX<sub>2-3</sub>S | RSpSDQS |

### Expression profile of phototropin and associated signalling proteins in wild type and CRISPR-Cas phototropin knockouts of *C. reinhardtii*

To further investigate phototropin’s interaction with other signaling proteins, qRT-PCR was used to compare the expression levels of photoreceptor genes (LOV-HK1, LOV-HK2, and COP6) between wild-type and phototropin knockout strains. Under prolonged blue light exposure, the expression of LOV-HK1 and LOV-HK2 decreased in wild-type cells, while COP6 remained elevated, potentially due to its role in high-light conditions (Fig. 9a). In contrast, the phototropin knockouts ΔPhot-B5 and ΔPhot-C41 (referred to as B5 and C4 in Fig. 9a) exhibited elevated expression of LOV-HK1 and LOV-HK2, suggesting their possible role in compensating phototropin function in blue light responses. This indicates the role of LOV-histidine kinases with phototropin in regulating blue light mediated responses in *C. reinhardtii*.

**Fig. 9.**
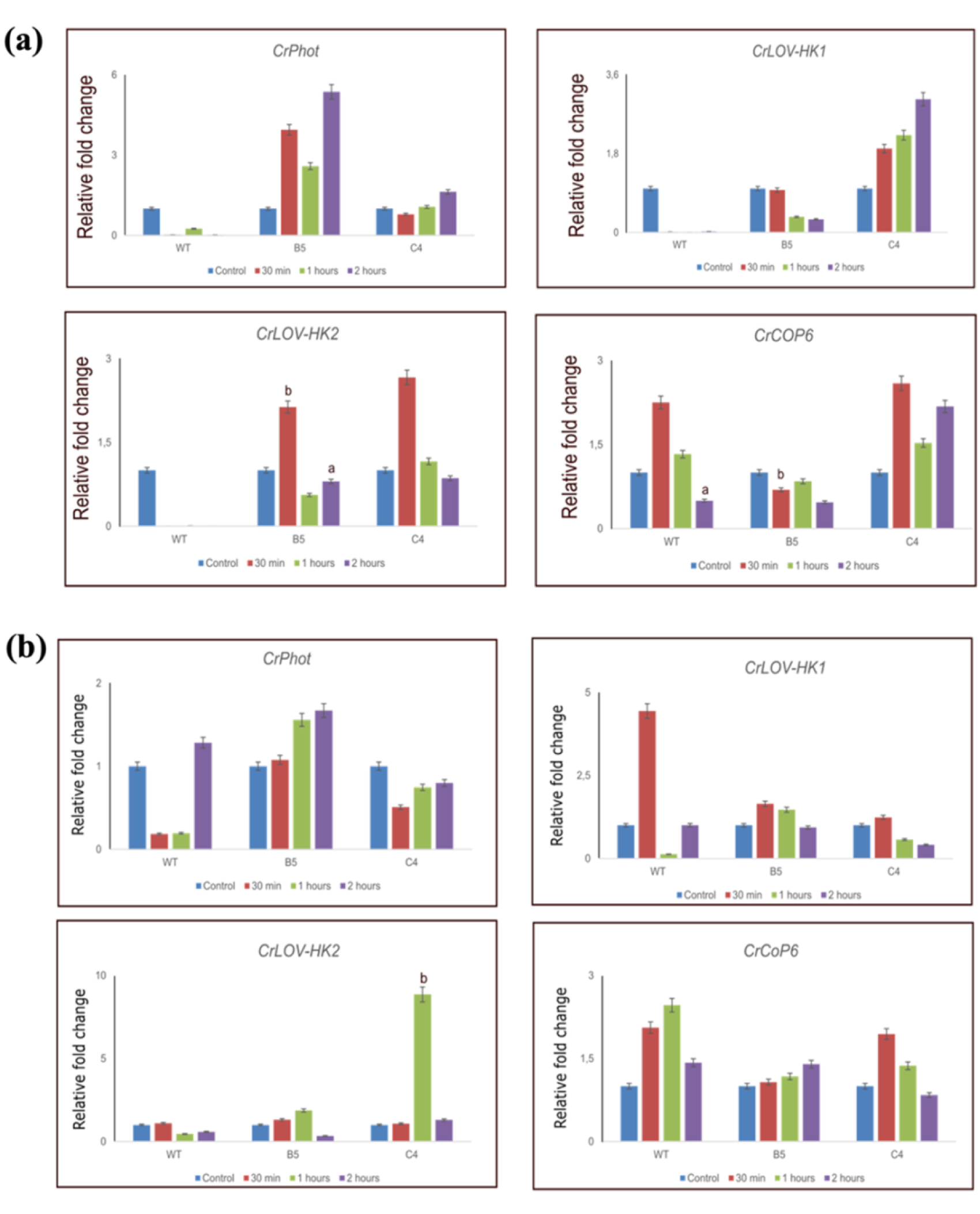
Gene expression profiles in light and dark exposed wild-type and phototropin mutants of *Chlamydomonas reinhardtii*. Alteration in gene expression of phototropin interacting genes like LOV histidine kinases (LOV-HK1, LOV-HK2) and COP6 was observed in blue light exposure and different light intensity. Expression profile of Phot, LOV-HK1, LOV-HK2, COP6 in (a) Light exposed samples (b) Dark exposed samples of wild type (CC125) & phototropin CRISPR-Cas generated mutants (B5: CC-125, ΔPhot-B5 & C4: CC-125, ΔPhot-C4-1) of *C. reinhardtii*, before and after blue light treatment (control samples are taken before blue light exposure). Statistical significance was determined using Student’s t-test: (b) *P* < 0.05, and (a) *P* < 0.005, compared to the control. The expression of all the genes were normalized with the expression of CBLP.

In dark exposed cells of phototropin knockout strains particularly, ΔPhot-C41 exhibited disrupted expression of these photoreceptor(s), particularly COP6 and LOV-HK1, indicating phototropin plays a key role in regulating the expression of other photoreceptors under varying light conditions (Fig. 9b).

Further, gene expression of photoreceptors (CrPhot, LOV-HK1, LOV-HK2, ChR1 and ChR2) were assessed in wild type strain of *C. reinhardtii* in different light conditions. Expression of CrPhot and ChR1 were elevated in red light whereas, LOV-HK1, LOV-HK2 and ChR2 expression were increased in UV light (Fig. S4).

### Role of phototropin and associated photoreceptor genes in gametogenesis in *Chlamydomonas reinhardtii*

It is known that phototropin plays a vital role in the sexual life cycle of *Chlamydomonas reinhardtii*, particularly during gametogenesis and gamete fusion [4]. To further investigate the involvement of phototropin and associated photoreceptor genes in gametogenesis, we examined the expression of several marker genes, including FUS1 (fusion marker), MID (minus dominance marker), GSP1 (plus gamete-specific marker), and GSM1 (minus gamete-specific marker). These genes are key indicators of different stages in the mating process, including gamete formation and zygote development [41–43]. The expression of phototropin was in corroboration with previous study [4].

Additionally, increased expression of photoreceptors, including channelrhodopsins (ChR1, ChR2) and 14-3-3 proteins, was observed, indicating potential interactions with phototropin. Similarly, LOV histidine kinases (LOV-HK1, LOV-HK2) also exhibited elevated expression levels (Fig. 10). These findings suggest that these proteins may participate in phototropin-mediated signaling pathways during the sexual life cycle.

**Fig. 10.**
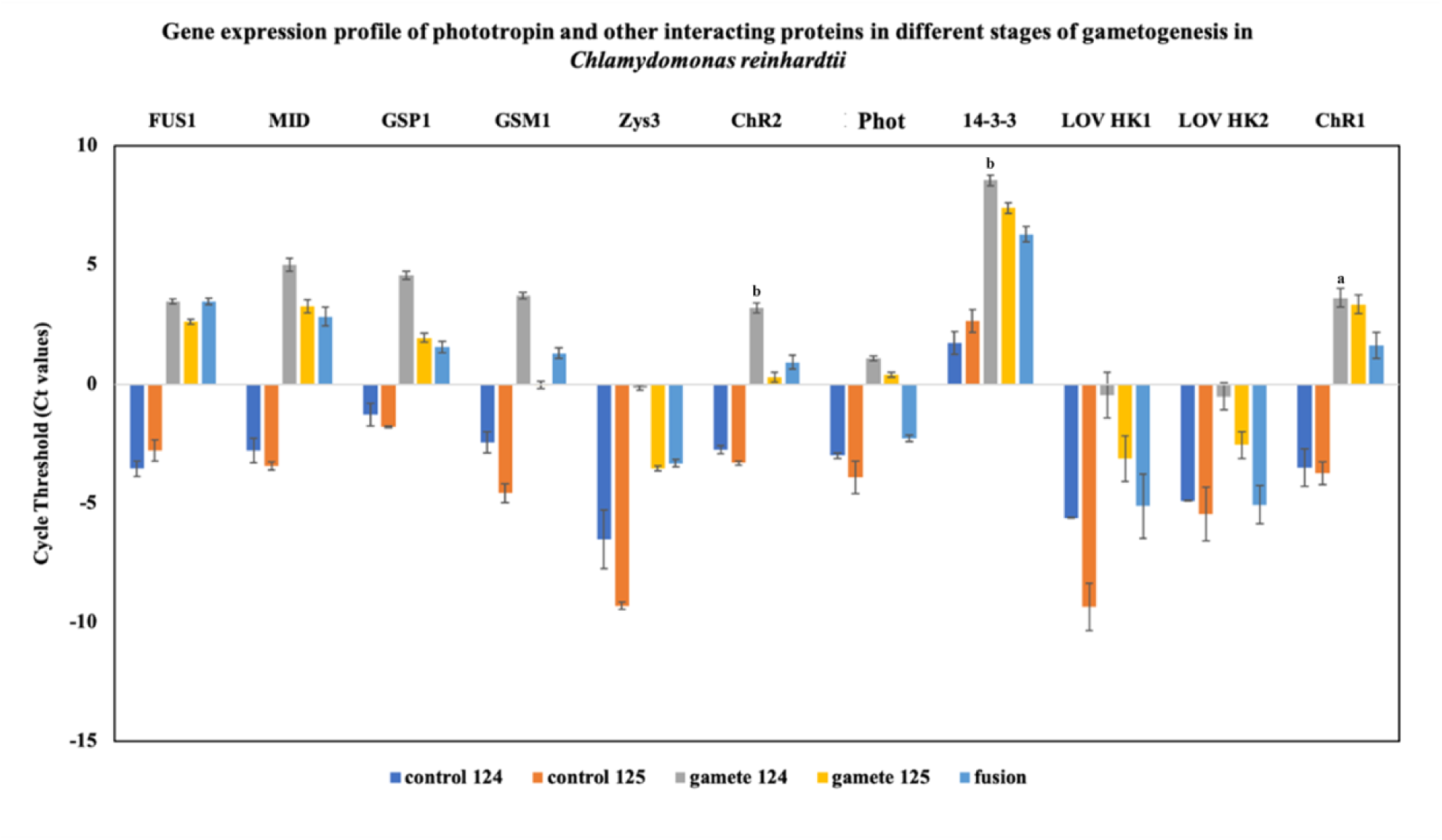
Gene expression profile of phototropin and other interacting proteins in different stages of gametogenesis in *Chlamydomonas reinhardtii*. The results shows that phototropin governs the gametogenesis and zygote development in coordination with other photoreceptor(s) genes (ChR1, ChR2, LOV-HK1, LOV-HK2) and signaling proteins (14-3-3). Different reporter genes (FUS1, MID, GSP1, GSM1, Zys3) specified for different stages of gametogenesis were used for studying expression profile. All genes were normalized with the expression of CBLP under particular control prior to the experimental control. Data represented are mean of three technical replicates and 3 biological replicates. Standard error bar represents the mean +/- SD. Statistical significance was performed using Student’s t-test: (b) *P* < 0.05, and (a) *P* < 0.005, compared to the control. ChR1, Channelrhodopsin 1; ChR2, Channerhodopsin 2; LOV-HK, LOV histidine kinase; Phot, Phototropin; CBLP, Beta subunit-like polypeptide; FUS1, fusion marker; MID, minus dominance marker; GSP1, plus gamete-specific marker; and GSM1, minus gamete-specific marker.

### Phototropin-regulated photomotility in *Chlamydomonas reinhardtii*

Phototropin plays an essential role in controlling photomotility in *Chlamydomonas reinhardtii* [3]. To elucidate the influence of phototropin on photomotility under blue light, wild type (CC- 125) and phototropin knockout strains (ΔPhot-B5 and ΔPhot-C4-1) were examined under dark and light-adapted conditions. Both knockout strains exhibited significantly reduced photomotility compared to the wild-type strain under all tested conditions (Fig. S5a). Upon blue light exposure (intensity ∼390-405 lux) for 30 and 120 minutes, partial recovery of photomotility was observed in the knockout strains, with the most pronounced response occurring at 120 minutes. Interestingly, light-adapted cells showed reduced photomotility compared to their dark-adapted counterparts, likely due to prior light saturation (Fig. S5b and c). Notably, the ΔPhot-B5 strain demonstrated minimal phototactic response, whereas the ΔPhot-C4-1 strain maintained some level of phototaxis, which further increased upon blue light exposure at the 120 minutes (Fig. S5b). Additionally, as phototropin plays a critical role in photosynthesis, the cellular growth of *Chlamydomonas* was monitored every 24 hours to generate growth curves for both the wild-type and phototropin mutant strains of *C. reinhardtii*. The deletion of the CrPhot gene significantly affected the growth of the mutant strains (CC- 125, ΔPhot-B5, and CC-125, ΔPhot-C4-1) (Fig. S6).

## Discussion

In the current study, we focussed on understanding the molecular mechanisms underlying phototropin localization, trafficking, and interactions in *Chlamydomonas reinhardtii* that can provide valuable insights into its role in mediating light-induced physiological processes.

Phototropin’s localization in the eyespot and flagella suggests its crucial function in photoreception and light signaling. The roles of these organelles in phototaxis and photosynthesis highlight phototropin’s integral position in coordinating light-mediated cellular responses [3]. Hence, altered or perturbed localization of phototropin can impair the normal photophysiological functioning in *C. reinhardtii*. The current research investigates the mechanistic basis of phototropin trafficking in *C. reinhardtii* using different IFT mutants. The immunolocalization results show that phototropin trafficking in different cellular location is intricately dependent on the intraflagellar transport (IFT) machinery. The findings from IFT52 and IFT88 mutants revealed that both IFT52 and IFT88 are critical for the anterograde transport of phototropin and flagellar assembly, consistent with their known roles in facilitating cargo movement along the flagellum [27,28]. Phototropin mislocalization in IFT52 (CC-477) and IFT88 (CC-3943) mutants, despite normal synthesis, suggests that these proteins are critical for its trafficking. Moreover, degradation of mislocalized phototropin indicates that accurate trafficking is required for its stabilization and function within the cell. Furthermore, accumulation of phototropin at the flagellar tip in IFT172 mutants demonstrates its role in switching of phototropin from anterograde to retrograde transport. Thus, highlighting the involvement of IFT components in phototropin recycling. This finding is supported by studies on retrograde IFT’s role in regulating flagellar protein turnover [34]. In IFT139 mutants, abnormal phototropin distribution near the basal body and eyespot emphasizes the critical function of the retrograde machinery in modulating photoreceptor localization and stability. The accumulation of phototropin at the flagellar tip in kinesin-2 and dynein double mutant strain CC-4424 suggests a disruption in its recycling, emphasizing the role of intraflagellar transport (IFT) machinery in maintaining phototropin’s dynamic localization and function. Specifically, kinesin-2, which drives anterograde transport, and dynein, responsible for retrograde transport, are both indispensable for the efficient trafficking of phototropin within flagella [34]. The CEP290 mutant strain (CC-4374), exhibits significantly reduced flagellar length, and an absence of phototropin near the basal body, likely due to disrupted IFT dynamics. Previous studies have established the importance of CEP290 in maintaining the integrity of IFT complexes [36], and our immunolocalization results suggest the essential role of CEP290 in ensuring phototropin stability and cellular localization. These findings corroborates with previous studies demonstrating the essential role of IFT in transporting photoreceptors and maintaining their functional distribution within flagella [44].

Our interactome analysis uncovered a complex network of interactions between phototropin and other photoreceptors, such as Channelrhodopsins (ChR1, ChR2), LOV-histidine kinases (LOV-HK1, LOV-HK2) and signalling protein such as 14-3-3 protein. Reduced expression of ChR1 and 14-3-3 in phototropin knockout strains suggest that phototropin modulates these proteins through a signaling cascade, a mechanism that may be conserved across species [45]. The presence of 14-3-3 binding motifs in phototropin and its homologs aligns with the interaction patterns observed in *Arabidopsis*, where phototropin regulates stomatal opening through 14-3-3 proteins [40]. These results suggest a broader regulatory role for phototropin in photoreceptor signaling networks across diverse species.

The role of phototropin in photomotility is well established, and this was further evidenced by phototropin knockout mutants, which displayed impaired motility under both light and dark conditions. Interestingly, partial recovery of phototaxis in knockout strains after prolonged blue light exposure suggests that other photoreceptors, such as ChR1 [3,46] and some unknown signaling proteins may compensate for the loss of phototropin function, albeit to a limited extent. The differential responses observed between the ΔPhot-B5 and ΔPhot-C4-1 strains further highlight the complexity of light perception and response mechanisms in *C. reinhardtii*. In *C. reinhardtii*, phototropin influences signalling cascades which controls the flagellar movement. The mislocalization of phototropin might disrupt the signalling between photoreceptors and flagella, consequently, perturbing phototaxis and other ciliary-mediated behaviour. Additionally, the growth defects observed in phototropin mutants likely reflect disruptions in photosynthesis, consistent with previous studies linking phototropin to photosynthetic regulation and photoprotection [47]. Furthermore, our interactome analysis indicated potential direct or indirect interactions between phototropin and other photoreceptors, suggesting a coordinated network that governs gametogenesis and zygote development. These data support the hypothesis that phototropin, through its involvement in photoreceptor signaling, plays a central role in modulating light-induced developmental processes in *C. reinhardtii* [48]. Ciliary targeting sequence (CTS) (Table S4) and post-translational modification (PTM) (Table S5) analysis of phototropin further reveals the adaptability of these proteins in response to environmental light cues. The extensive PTM patterns identified in this study highlight the dynamic regulation of photoreceptors and underscore the importance of PTMs in fine-tuning of light perception and signaling. Such modifications may enhance the ability of *C. reinhardtii* to respond to fluctuating environmental conditions, ultimately optimizing its phototrophic lifestyle.

In conclusion, this study showed the critical roles of IFT machinery, photoreceptor interactions, and PTMs in regulating phototropin localization, stability, and function in *C. reinhardtii*. Future research should focus on elucidating the molecular details of these regulatory networks, particularly the cross-talk between IFT components, photoreceptor signaling pathways, and PTMs. Such studies will be pivotal in advancing our understanding of phototropin’s multifaceted role in environmental sensing and cellular adaptation in algae.

## Abbreviations

Phot, Phototropin; LOV, Light-Oxygen-Voltage; STK, Serine-threonine kinase; ChR, Channelrhodospin(s); HK, Histidine kinase; CBL, Calcineurin B-like proteins; CBLP, Beta subunit-like polypeptide; PTM, Post-translational modifications; CEP290, Centrosome-associated protein; CTS, Ciliary targeting sequence; CAM, Calmodulin; ARF, ADP ribosylation factor; min, minutes; h: hour; Act.: Acetylated: SF: Soluble fraction; TCL: Total cell lysate; IFT: Intraflagellar transport; PTM: Post-translational modification.

## Author Contributions

S.K. conceived the project. S.S. designed experiments under the guidance of S.K. and drafted the manuscript with contributions from S.K., R.S., K.S., and S.K.S. S.S. conducted immunofluorescence, immunoblotting, qRT-PCR, photomotility, and bioinformatics experiments for the publication. K.S. contributed to bioinformatics, qRT-PCR, immunoblotting and immunofluorescence. R.S. contributed immunoblotting and image processing. S.K.S. contributed to the qRT-PCR analysis. All authors reviewed and approved the final manuscript.

## Data availability

Data will be made available on request.

## Acknowledgements

We would like to express our gratitude to Prof. Peter Hegemann from Humboldt University, Germany, for generously providing us with the phototropin mutant strains (Phot-B5 and Phot- C4-1). S.S. is highly thankful to CSIR for providing financial assistance. RS acknowledge DBT-RA program (DBT-RA/2022/July/N/2560). We gratefully acknowledge S.K. lab members for their suggestions. The authors acknowledge Dr. Neetu Singh and the Advanced Instrumentation Research Facility (AIRF), Jawaharlal Nehru University.

