## Supplementary material for "Phototropin localization and interactions regulates photophysiological processes in *Chlamydomonas reinhardtii*": Supplementary information.docx

**Material and methods**

**Post-translational modification (PTMs) prediction**

Predictions of various PTMs were carried out using specialized software. SUMOylation sites were predicted via GPS SUMO version 2.0 (http://sumosp.biocuckoo.org/citation.php) [8] and JASSA version 4 (<http://www.jassa.fr/index.php>.) [8]. Lipid modifications, specifically palmitoylation and myristoylation, were assessed using the GPS-lipid server version 1.0 (<http://lipid.biocuckoo.org/>) [9] and CSS-PALM version 4.0 (<http://csspalm.biocuckoo.org/>) [10], respectively, along with the Expasy-Myristoylator tool (<https://web.expasy.org/myristoylator/>) [11]. Acetylation sites (N-acetylation and internal lysine acetylation) were predicted using NetAcet 1.0 (<http://www.cbs.dtu.dk/services/NetAcet/>) [12] and GPS-PAIL 2.0 (<http://pail.biocuckoo.org/>) [13,14]. Nuclear localization and export signals were identified using NLStradamus (<http://www.moseslab.csb.utoronto.ca/NLStradamus/>) [15] and LocNES (<http://prodata.swmed.edu/LocNES/LocNES.php>) [16] or NetNES 1.1 (<http://www.cbs.dtu.dk/services/NetNES/>) [17], respectively. Initiator methionine cleavage and Nα-terminal acetylation were analyzed using the Terminus server (<http://terminus.unige.ch/>) [18]. Glycation sites were predicted with NetNGlyc 1.0 (<http://www.cbs.dtu.dk/services/NetNGlyc/>) [19]. Phosphorylation and ubiquitinylation sites were predicted using GPS version 5.0 (<http://gps.biocuckoo.cn/wsresult.php>) [20] or NetPhos 3.1 server (<http://www.cbs.dtu.dk/services/NetPhos/>) [21,22] and BDM-PUB (<http://bdmpub.biocuckoo.org/prediction.php>), respectively.

**Phototaxis assay:**

In the phototaxis assay, petri dishes with uniformly distributed algal populations were illuminated from one side with white light (~1500-2000 lux) for 10 minutes to assess phototactic behavior. After that image were captured manually using the camera. A modified Petri dish assay was used to compare phototactic responses between *Chlamydomonas reinhardtii* wild type and phototropin mutants. For the experiment 4 hours light and 14 hours dark-adapted algal cells were used. Petridishes with equal number of algal cells, were exposed to blue light (~400 lux) for 30 and 120-minute intervals in petridishes. Phototaxis was assessed by capturing an image after 30 and 120 minutes manually with camera and quantifying movement towards the light source by comparing images before and after exposure. Cells without blue light exposure were used as control for comparison of cellular photomotility response.

**Supplementary figures:**

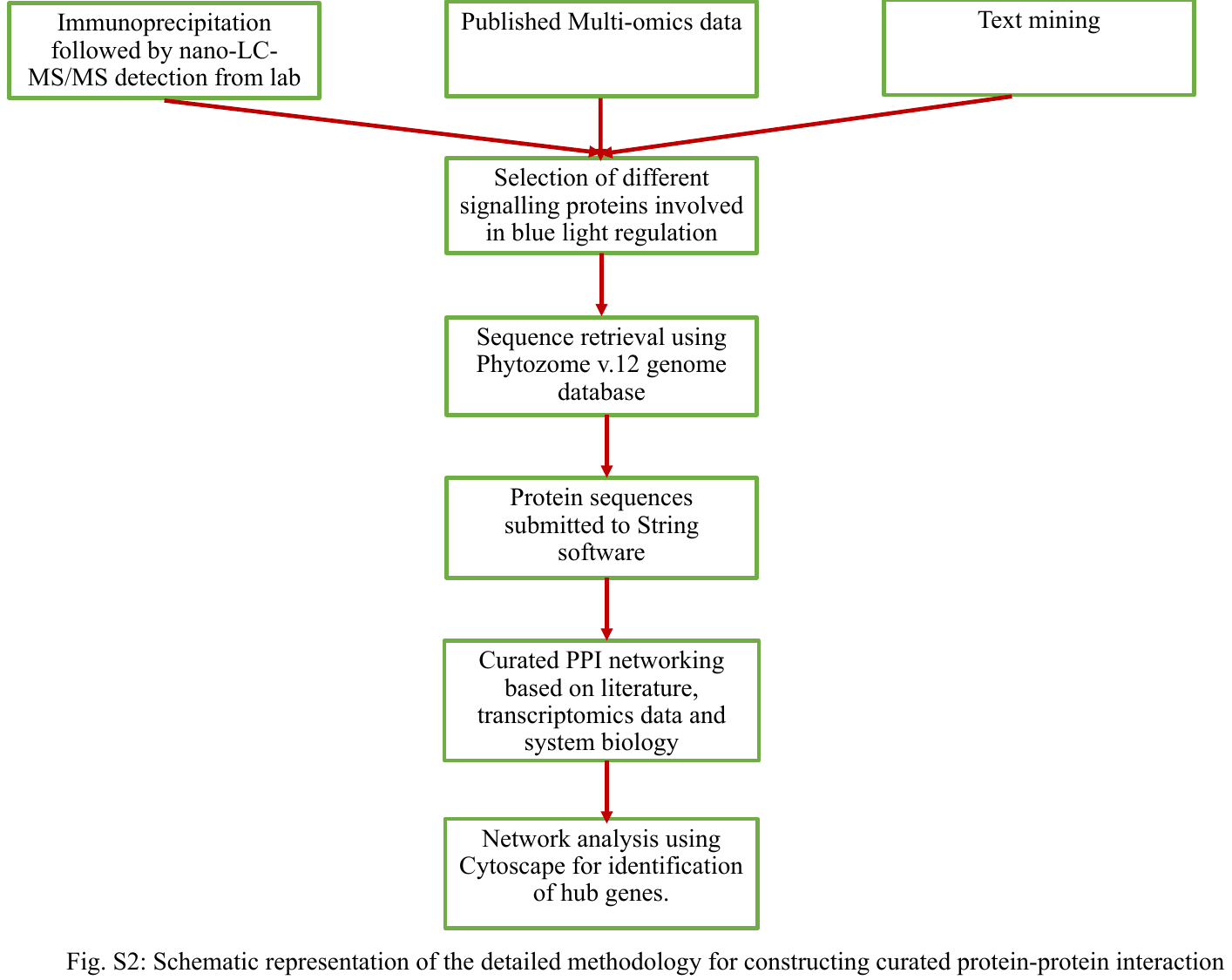

**Fig. S1. Schematic representation of the detailed methodology for constructing curated protein-protein interaction.**

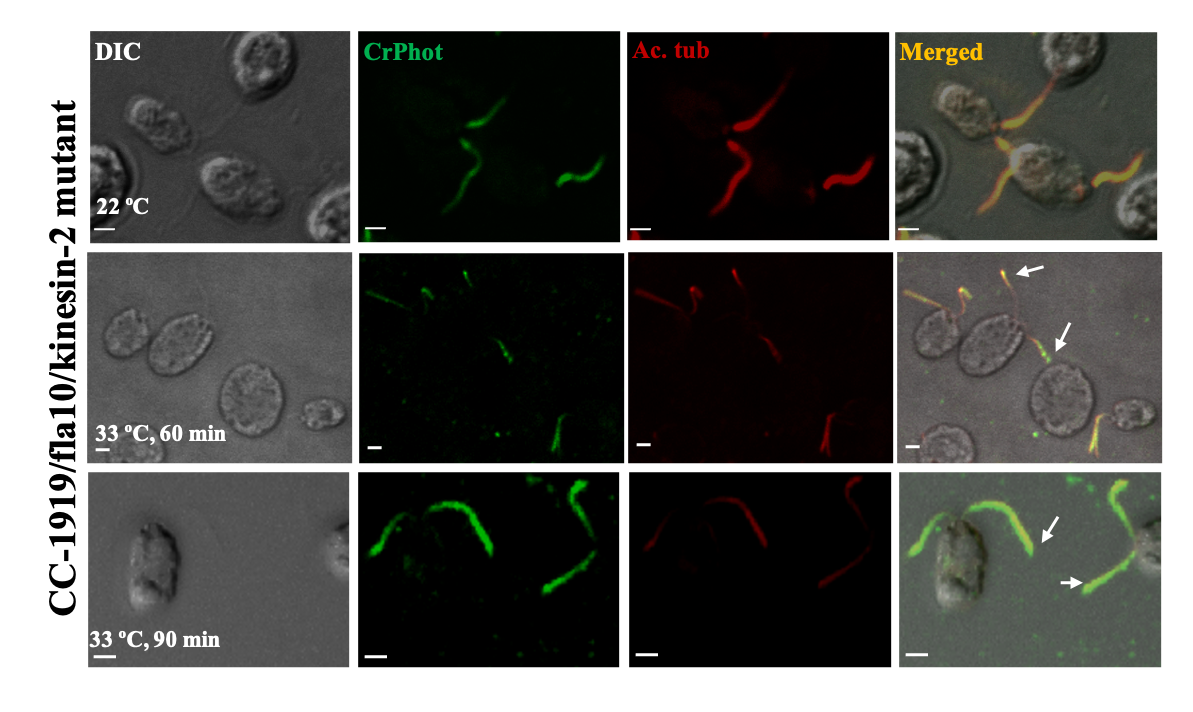

**Fig. S2. Immunolocalization of CrPhot in Kinesin motor mutant strain (CC-1919) of *C. reinhardtii* cells.**

The accumulation of phototropin was observed at the flagellar tip. The first panel represents the DIC image of algal cells. The second panel represents the CrPhot signal (green) with primary antibody anti-cLOV (dilution-1:250) and secondary antibody anti-goat IgG Alexa-488 (dilution-1:1000). The third panel represents the signal for acetylated tubulin (red) with primary antibody anti-act. tubulin (dilution-1:1000) and secondary antibody anti-mouse IgG Alexa-647 conjugated (dilution-1:1000). The fourth panel represents the overlay (merged) of the first three panels. Scale bar = 2 μm. The arrow indicates accumulation of phototropin.

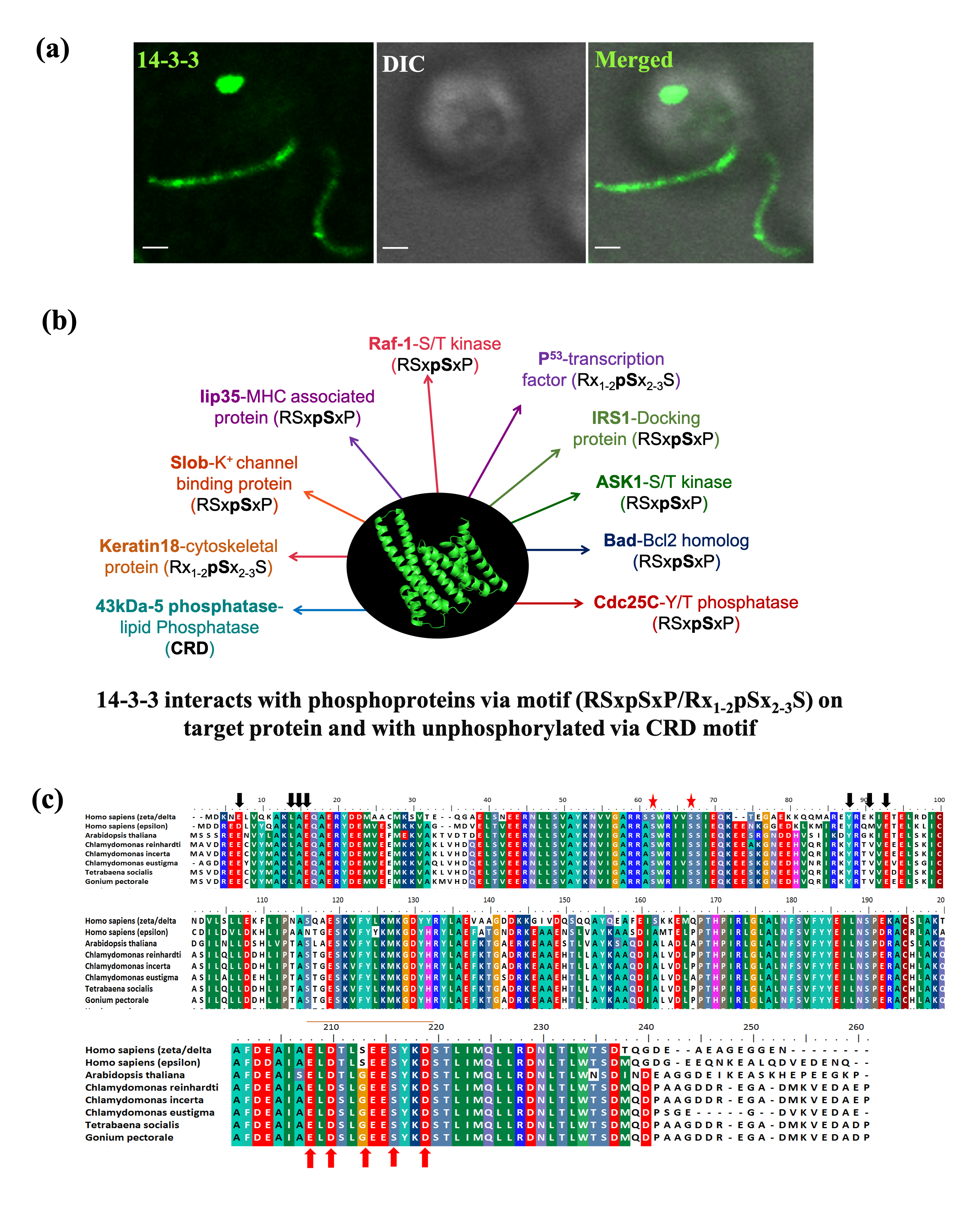

**Fig. S3. Differential interactions of 14-3-3 protein with different unphosphorylated and phosphorylated target protein molecules.**

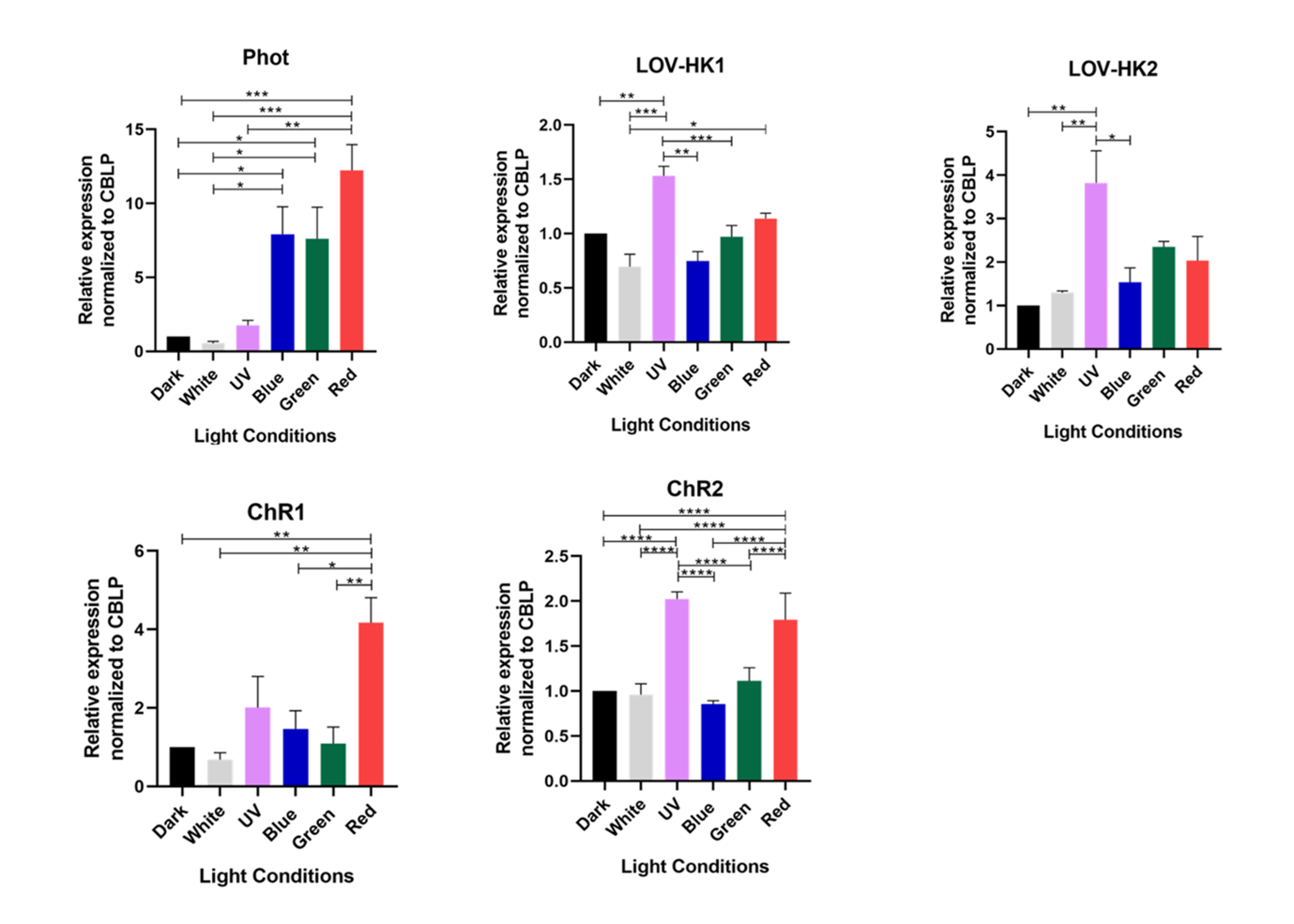

**Fig. S4. Gene expression profile of phototropin and interacting light sensing proteins in *Chlamydomonas reinhardtii* under varying light conditions.**

Altered expression of photoreceptor(s) was observed under different light conditions. Expression profile of *Chlamydomonas* different genes like phototropin (Phot), LOV-histidine kinase(s) (LOV-HK1, LOV-HK2), channelrhodopsins (ChR1, ChR2) in different light conditions. Control samples have been taken in dark and mRNA levels of all genes were normalized with CBLP gene under similar condition. Data represented are the mean of three biological replicates under a particular condition. The standard error bar represents the mean +/- SD of fold change. CBLP, Beta subunit-like polypeptide. The samples were exposed for 2 hours after the end of dark cycle at ~400 lux light intensity.

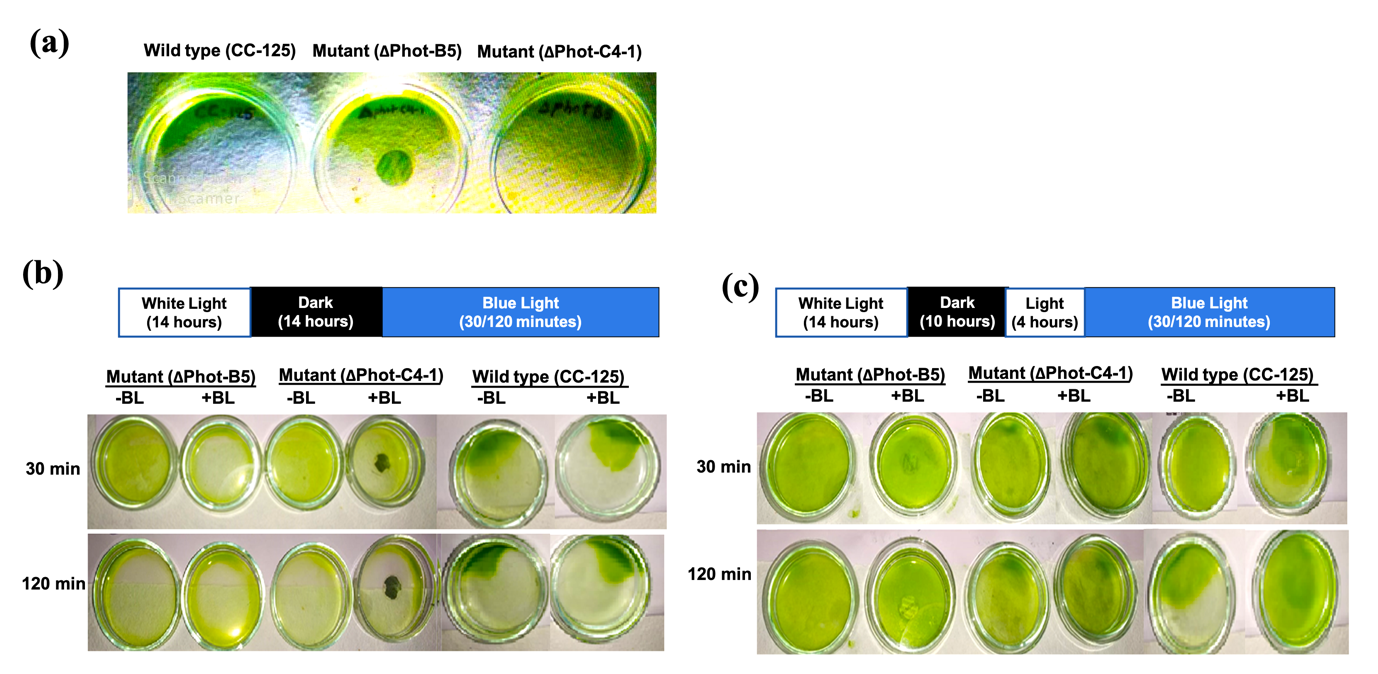

**Fig. S5. Phototropin-dependent alterations in phototactic behavior of *Chlamydomonas reinhardtii* under blue light exposure.**

(a) Comparison of phototaxis in wild type (CC-125) and phototropin knockout (ΔPhot B-5 and ΔPhot-C4-1) strains under 10 minutes of white light. (b) Phototactic response of dark-adapted cells and (c) light-adapted cells in wild type (CC-125) and phototropin knockout mutants (ΔPhot-B5 and ΔPhot-C4-1) in blue light (+BL: with blue light; +BL: without blue light) exposure of 30 and 120 minutes.

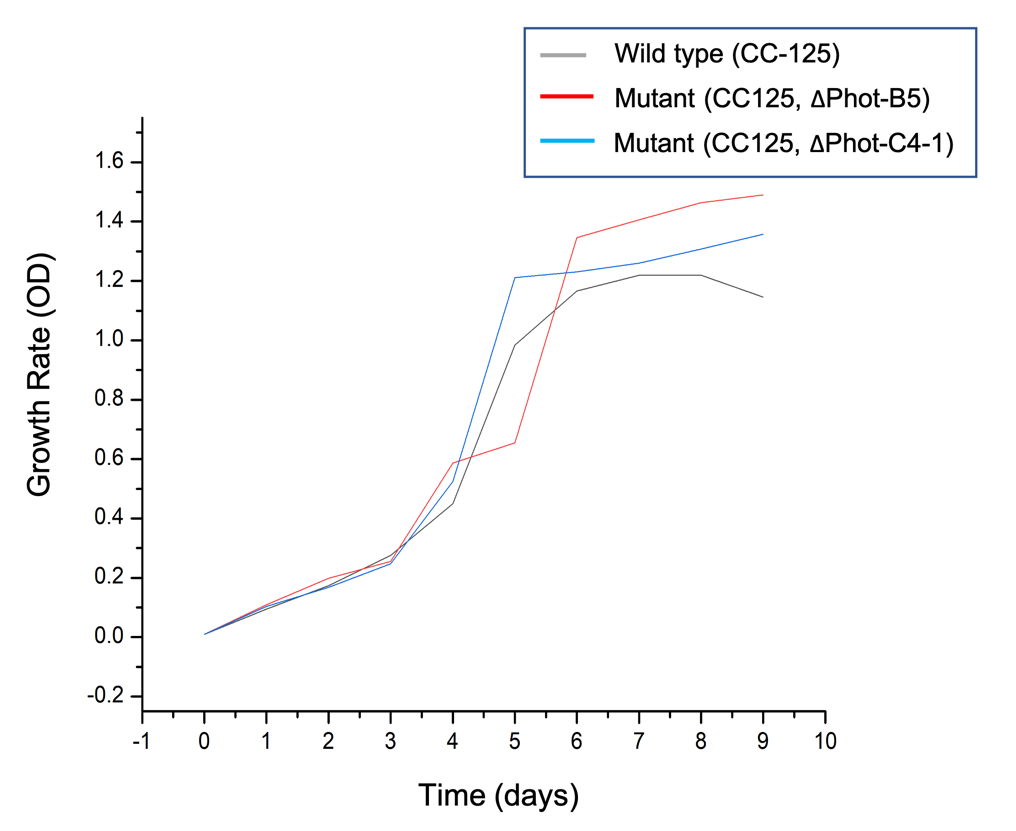

**Fig. S6. Growth curves of wild-type (*CC-125*) and phototropin mutant strains (ΔPhot-B5 and ΔPhot-C4-1) of *Chlamydomonas reinhardtii*.** Growth was monitored under standard conditions to compare the proliferation rates between wild type and phototropin-deficient strains.

**Supplementary Tables:**

**Table S1. Accession numbers of phototropin and associated interacting protein sequences identified in Chlamydomonas reinhardtii.**

| ***Gene name*** | ***Sequence identifier*** |
| --- | --- |
| Phototropin | Cre03.g199000 |
| LOV-HK1 | Cre13.g571200 |
| LOV-HK2 | Cre02.g079750 |
| 14-3-3 | Cre12.g559250 |
| ChR1 | Cre14.g611300 |
| ChR2 | Cre02.g085257 |
| COP6 | Cre11.g467678 |

**Table S2. List of *Chlamydomonas* strains used in the study.**

| **Strain** | **Growth temperature** | **Remarks** |
| --- | --- | --- |
| CC-124 137c mt- | 22˚C | Wild type |
| CC-125 137c mt+ | 22 ˚C | Wild type |
| CC-1396 fla8 mt- | Permissive 22 ˚C  Non-permissive 33 ˚C | Kinesin-2 motor protein mutant |
| CC-1919 fla10 | Permissive 22 ˚C  Non-permissive 33 ˚C | Kinesin-2 motor protein mutant |
| CC-3937 fla14-1: NIT1 mt+ | 22 ˚C | Dynein light chain |
| CC-4424 fla10, dhc1b-3 | Permissive 22 ˚C  Non- permissive 33 ˚C | Kinesin-2 motor/dynein heavy chain double mutant |
| CC-477 | 22 ˚C | Ift52 (complex b) |
| CC-3943 | 22 ˚C | Ift88 (complex b) |
| CC-1920 fla11 mt- | Permissive 22 ˚C  Non-permissive 33 ˚C | Ift172 (complex b) |
| CC-4374 cep290 mt+ | 22 ˚C | CEP290 (Centrosome associated protein) |
| CC-3863 fla17-1 mt+ | Permissive 22 ˚C  Non- permissive 33 ˚C | IFT139 (Complex A) |

**Table S3. List of primers used for qRT-PCR analysis of various genes in *Chlamydomonas reinhardtii*.**

| ***Primer name*** | ***Primer sequence (5’-3’)*** |
| --- | --- |
| CrFUS1_qFwd | CGGCAGCGAAGTTCCCATACG |
| CrFUS1_qRev | CTTGTGGCAAGCCGCTTGCAG |
| CrMID_qFwd | GACTCTGGTGCGGGCTCATCG |
| CrMID_qRev | TCGTCCGTGTCGAGGAGCATC |
| CrGSP1_qFwd | CAGCGACAAACGACCGTGCGG |
| CrGSP1_qRev | AACTGCTGCGCCATCCATCGC |
| CrGSM1_qFwd | GCGCACGAGACACCAGGACA |
| CrGSM1_qRev | CAGCATGTGAAGGCTGCTGCC |
| CrZys3_qFwd | TGCGCGCCTCTCCACCGAAC |
| CrZys3_qRev | GACACGCAGCCTAGCGCGAA |
| CrChR1_qFwd | GCTTGAGCACACCGCTGAGC |
| CrChR1_qRev | GCGTCACTGCCAGTCGAGGC |
| CrChR2_qFwd | CATACTTGACATCTGTCGCC |
| CrChR2_qRev | ACGACTACTGGGTTCGTTAC |
| CrCOP6_qFwd | CACCGGCTTCTTTGTCTACATAACTG |
| CrCOP6_qRev | AGTGTTGGCACCTGGGCCCG |
| Cr14-3-3_qFwd | CTGAGATGGCGGTCGACCGGGAG |
| Cr14-3-3_qRev | CTTGGCCACCTTCTTCATCTCCTCC |
| CrCBL1_qFwd | CTGTAGCCTGCAAACAGTAG |
| CrCBL1_qRev | TGCCCGTGTTGTTGGCAAGG |
| CrCBL2_qFwd | AGCTCGTGCAGCTCTACCAT |
| CrCBL2_qRev | TTGTACAGCGCCTCTATCTC |
| CrCBLP_qFwd | CTTCTCGCCCATGACCAC |
| CrCBLP_qRev | CCCACCAGGTTGTTCTTCAG |
| CrPhot_qFwd | ACAGCCATGGCAGGGGTGCCAG |
| CrPhot_qRev | ACAGTCCGGAAGCGTGGCATCT |
| CrLOV-HK1_qFwd | CGAGCAGAATGGACGCTGCTCT |
| CrLOV-HK1_qRev | CGAGGTAGCCCTCTGCTTGCTG |
| CrLOV-HK2_qFwd | AACAAAATGGACCGGAGGGAG |
| CrLOV-HK2_qRev | GCCTGTTCGGCAATTCCAGTAA |

**Table S4. Phototropin ciliary targeting sequence (CTS) in *Chlamydomonas reinhardtii.***

| ***CTS motif*** | ***Sequence^#^*** |
| --- | --- |
| AXXXQ motif [49] | MAGVP**APASQ**LTKVLAG |
| FR motif [50] | DGKMKLMH**FR**RVKQLGAG, LYGTTP**FR**GARRDE |
| VxPx motif [51][49][52] | TPFWNLLT**VTPI**KTPDGRV, TIAVEKA**E**G**VEPG**QASAVAAAA, GYVYRD**LKPE**NILLHHTG, GTEEY**LAPE**VINAAG, FWNMFT**LAPM**RDQDGHARF |

**#**Residue highlighted in bold indicates the conserved amino acid residue in the sequence.

**Table S5. Different post-translational modifications in phototropin of *C. reinhardtii*.**

| ***Modification Type*** | ***Position*** | ***Consensus Sequence/Motif*** | ***Details*** |
| --- | --- | --- | --- |
| SUMOylation | K143, K502, K531 | SGVPLLVKYDHRLRD, SQPKKRLKEEHVRFY, GYVYRDLKPENILLH | SUMOylation at specific lysine residues |
|  | K384, K398 | MRRKPHKADDKAYQALL, ALLQLQERDGKMKLMHF | SUMOylation at inverted consensus sites |
| Phosphorylation | S581 | SPKKSSSKS | Kinases: Unspecified, PKC, PKG; Scores: 0.979, 0.722, 0.587 |
|  | S577 | SAPKSPKKS | Kinases: Unspecified, cdk5, GSK3; Scores: 0.997, 0.713, 0.647, 0.535 |
|  | S582 | PKKSSSKSG | Kinases: Unspecified, PKC; Scores: 0.993, 0.821 |
|  | S585 | SSSKSGGSS | Kinases: Unspecified, PKC; Scores: 0.994, 0.688 |
|  | S679 | KPAVSEECR | Kinase: Unspecified; Score: 0.994 |
| Ubiquitination | 576 | AAGGSAPKSPKKSSS | Ubiquitination at lysine residues in specified motifs |
|  | 579 | GSAPKSPKKSSSKSG | |
|  | 580 | SAPKSPKKSSSKSGG | |
|  | 584 | SPKKSSSKSGGSSSG | |
|  | 735 | YVPRRASKAAGGSST | |
| Lipid Modification | 216 | ERIQQNFCISDPTLP | S-Palmitoylation at Cluster B |
| Acetylation | K187 | VQREK | Internal lysine acetylation |
|  | K380 | NKEKL |  |
|  | K735 | MRLKY |  |
|  | K580 | SKLRK |  |
|  | K584 | EKKPT |  |
| Nuclear Import Signal | 497-503 | PKKRLKE | |
|  | 558-589 | TPRIEKIGGAGAAGGSAPKSPKKSSSKSGGSS | |
| Nuclear Export Signal | 148-162 | LRDNVARTIVDDVTI | |
|  | 403-417 | HFRRVKQLGAGDVGL | |
| Glycation | 282 | NYTK | Glycation at specific motif |

**Table S6. Interactions predicted with phototropin and their UniProt IDs in *Chlamydomonas reinhardtii* using STRING software.**

| ***Protein Name*** | ***UniProt ID*** |
| --- | --- |
| COP3, Channelrhodopsin 1 | A8JAJ2 |
| COP4, Channelrhodopsin 2 | Q8RUT8 |
| ARF (Arl3), ADP ribosylation factor-like 3 | A8ISN6 |
| PHOT, Phototropin | Q8LPD9 |
| CAM1, Calmodulin | P04352 |
| FTT1, 14-3-3 protein | P52908 |
| COP5, Chlamyopsin 5 | A8I9V1 |
| COP6, Chlamyopsin 6 | A0A0D2M2L8 |
| NIT1, Nitrate reductase | A8J4P9 |
| SUOX1, Sulfite oxidase | A8JEP4 |
| PIN4, Peptidyl-prolyl cis-trans isomerase | A8J3E3 |
| Calci_B (CBL), Calcineurin-like phosphoesterase B | A8JHJ4 |
| PIN3, Peptidyl-prolyl cis-trans isomerase | A8IU62 |
| NCS6, Cytoplasmic tRNA 2-thiolation protein 1 | A8JF71 |
| CGL49, ARF/SAR superfamily GTP binding protein | A8ILA3 |
| URM1, Ubiquitin-related modifier 1 homolog | A8IC48 |
| TST, Sulfur-transferase | A0A2K3D1J4 |
| APG8, Autophagy-related protein | A8JB85 |
| MCM5, DNA helicase | A8HPZ4 |
| EB1, Microtubule plus-end binding protein | Q84TR5 |
| CAV2, Voltage-gated Ca2+ channel, alpha subunit | A8JF90 |
| FAP259, Flagellar associated protein | A8ITN7 |
| BLD1 (IFT52), Intraflagellar transport protein 52 | Q946G4 |
| FAP60, Flagellar associated protein | A8HWB9 |
| IFT140, Intraflagellar transport protein 140 | A8J4D9 |
| BBS1, Bardet-Biedl syndrome 1 protein | A8JEA1 |
| IFT20, Intraflagellar transport protein 20 | Q8LLV9 |
| IFT172, Intraflagellar transport protein 172 | Q5DM57 |
| FLA8, Kinesin-like protein | A7LGV1 |
| FLA10, Kinesin-like protein | P46869 |
| IFT81, Intraflagellar transport protein 81 | Q68RJ5 |
| DHC2, Dynein heavy chain 2 | A8J063 |
| IFT88, Intraflagellar transport protein 88 | A8JCJ2 |
| FAP32 (IFT46), Intraflagellar transport protein 46 | A2T2X4 |
| LOV-HK1, LOV-histidine kinase 1 | A0A2K3CZT1 |
| LOV-HK1, LOV-histidine kinase 2 | A0A2K3E0G1 |
